# Differential Expression and Microsystem Physiology Reveal Predominant and Drug Reversible CFTR-Related Defects in Idiopathic Pancreatitis

**DOI:** 10.1101/2025.05.25.656006

**Authors:** Paba Edisuriya, Pramodha Liyanage, Jesun Lee, Anusha Jayesh Shetty, Hongyu Liu, Nicholas Tatonetti, Danielle Hutchings, Alvin J. Freeman, Jaimie D. Nathan, Appakalai N. Balamurugan, Stephen J. Pandol, Anil G. Jegga, Anjaparavanda P. Naren, Kavisha Arora

## Abstract

Pancreatitis is a potentially fatal and difficult to control exocrine-tissue defect with no FDA approved therapies. Variants of a chloride/bicarbonate transporter cystic fibrosis transmembrane conductance regulator (CFTR) pose multi-fold increased risk of pancreatitis accounting for up to 40% of the patients with idiopathic pancreatitis. However, the relationship between the duct-restricted CFTR-function and total exocrine tissue defect during pancreatitis remains less known and animal models do not translate well to human disease. To overcome this challenge, we developed a robust and highly durable iPSC-derived model system of pancreatic ductal tissues from an idiopathic pancreatitis patient with a common pancreatitis-associated *CFTR* variant. In the patient line termed PANx, we found deficient CFTR function and a distinct gene expression signature for ductal tissue pancreatitis marked by aberrant mucin production, inflammatory cytokines and cystic neoplasms. By applying clinically used CFTR-modulator drug ivacaftor, we observed a remarkable restoration of deficient CFTR-mediated fluid secretion as well as upto 40% reversal of the differential gene signature for PANx including the reduction in mucinous neoplasms and immunogenic cytokines such as IL-11, CCL20 and CXCL8. We further employed a microsystem device to model hyperamylasemia, a diagnostic feature of acute pancreatitis attack, due to a ductal reaction causing acinar injury. The key mucinous signature was validated in primary pancreatitis ductal tissues with a *CFTR* variant. Overall, we unraveled new layers of CFTR-related pathology in pancreatitis to help us better understand the early course of this debilitating condition. The test methods and model systems discovered in this study will significantly expedite the discovery of diagnostic and therapeutic tools for treating idiopathic pancreatitis. For the first time, we provided molecular and physiologic evidence supporting the benefit of CFTR modulator drug ivacaftor in human CFTR-related pancreatitis.

## Introduction

Acute (AP), recurrent acute (RAP), and chronic pancreatitis (CP) are potential life-threatening conditions and together remain one of the most common gastrointestinal-related cause of hospitalization in the United States^1–4^. Unabatable abdominal pain is the primary prompt for hospital admission for pancreatitis patients. The lack of any therapy in pancreatitis renders this condition extremely difficult to control. The impact of pancreatitis on patients’ quality of life is substantial, as they struggle with performing daily activities and suffer from the fear of social stigma, unemployment, and depression^5,6^. Post two–three decades into the diagnosis of CP, patients have dismal survival rate of 50%^2^. The cause of AP is sudden and could be extra-pancreatic while CP is long term and associated with pancreas scarring commonly caused by alcohol toxicity, RAP episodes, or susceptibility genes^1,7^. The less common causes of pancreatitis are included in M-ANNHEIM classification system^8^. There is known to be a disease continuum among AP, RAP, and CP, with more than 10% of patients with a first episode of AP and 36% of patients with RAP developing CP^9,10^.

Hereditary pancreatitis is caused by autosomal dominant mutations in the cationic trypsinogen gene (*PRSS1*) while other significant genetic susceptibilities are known including for haplotypic variants in cystic fibrosis transmembrane conductance regulator (*CFTR*) and serine protease inhibitor Kazal type 1 (*SPINK1*) genes^11,12^. Large epidemiological studies have demonstrated that carriers of *CFTR* variants are at a four-fold increased risk of developing idiopathic pancreatitis^13^. This has accounted for as many as 40% of pancreatitis patients carrying *CFTR* variants. Further *CFTR* variants increase the risk of patients undergoing total pancreatectomy with islet autotransplantation (TPIAT), which provides relief from pain but leads to long-term morbidities due to the lack of panceas-associated function^12–14^. Common pancreatitis causing insults, such as alcohol, smoking, and bile acids are also known to strongly inhibit CFTR function^11,15–17^. Thus a key factor between genetic and environmental pancreatitis etiologies is the perturbation of the CFTR protein which is largely expressed in the pancreatic ducts^11,15^. The precise mechanisms linking a lack of CFTR activity to the progression of pancreatitis including of the acinar in a genetically predisposed duct remain poorly understood. A significant limitation in the field is the absence of a relevant human-based model system to advance our understanding of pancreatitis pathophysiology. Human pancreases are scarce for research purposes, and primary tissues lack the necessary durability for long-term study^18,19^. Existing animal models, especially regarding CFTR-related pathophysiology, do not adequately replicate the characteristics and severity of human pancreatitis^20–22^. To overcome this limitation, we relied on induced pluripotent stem cells (iPSCs) technology to develop a human relevant model system of the pancreatic ductal tissue to study CFTR-related pancreatitis. Why study ductal tissue for pancreatitis? We know that ductal passage is central to the development of pancreatitis because of the structural or physiological conditions leading to ductal obstruction that causes insufficient flushing of the acini by intralobular ducts culminating into auto-digestion and damage of acinar cells by hyper-concetrated enzymes within the acini. Indeed, ductal ligation is a highly adopted animal model for pancreatitis^23,24^. Currently, there is no direct experimental evidence to support the involvement of ductal function in causing exocrine damage during pancreatitis and due to delayed diagnosis early disease patterns of pancreatitis in the ducts have been rarely identified. Further, it remains unknown whether a deficit in CFTR function leading to a dysfunctional duct could perpetuate this cycle of ductal-acinar damage, a condition that could possibly be ameliorated using CFTR-activating agents. As of today many of these agents are clinically approved and highly effective in the treatment of Cystic Fibrosis (CF)^25,26^. The possibility of that CFTR-agents are beneficial in the treatment of pancreatitis has only been explored in a limited number of studies which are largely observational^27–29^. Establishing CFTR dysfunction as a cause or contributor to idiopathic pancreatitis is critical to develop care plan for these patients and understand the systemic scope of their disease, and potentially allow their access to approved treatments.

By performing directed differentiation of induced pluripotent stem cells (iPSCs), we derived ductal tissues from a patient with idiopathic RAP containing a common pancreatitis *CFTR* variant V470M. This patient line also contains a mutation in CFTR modifier gene *SLC26A9*, a chloride/bicarbonate exchanger and no variants for acinar cell susceptibility genes *SPINK1* and *PRSS1*. Developing *de novo* ductal tissue from this line is ideal to study genetically-driven ductal tissue pathology in pancreatitis to study early disease patterns without acinar involvement or overt injury. Based on the comprehensive studies of iPSC-derived pancreatic cells, we identified CFTR dysfunction in ductal cells from RAP patient leading to hypermucinous state characterized by multifold increased expression of membrane-bound mucin MUC1 as well as secreted mucins normally unexpressed by ductal cells. We additionally identified a specialized mucin expressing ductal population as a characteristic of this patient line that could potentially be associated with serious sequential pancreatic morbidities including the development of hypermucinous neoplasia and pancreatic duct associated adenocarcinoma (PDAC). This state was remarkably reversible using the CFTR potentiator, ivacaftor. Indeed, a significant proportion of the differential expression state of patient cells including for the NF-κB pathway that was here identified in ductal cells, could be restored to the control levels upon ivacaftor treatment suggesting a signficant drug-reversible nature of pancreatitis pathology. Using a microsystem device carrying pancreatitis patient-derived acinar cells, we established hyperviscous ductal secretion as a driver of reaction for hyperamylasemia, a feature of acute pancreatitis attack. We hypothesized that a continuum of hypermucinous and hyperviscous ductal secretion, leads to chronic acini injury and the development of pancreatitis. Overall, we discover new aspects of pancreatitis pathology and model system utilization that would accerlerate the investigation of novel molecular drivers, therapeutic targets, and diagnostics for pancreatitis.

## Results

### Development of iPSC-derived human model of the exocrine pancreas containing ductal tissues that closely mimics primary culture signature and growth dynamics

Pancreatic exocrine and endocrine cells originate from a common precursor, namely the pancreatic progenitor (PP) cell^30,31^. PPs acquire a tripartite state with their tip domain to give rise to the acinar lineage, while a bipotent trunk domain forms the ductal lineage together with subsequently stratifying endocrine cell types^32,33^. We followed an earlier published protocol to capture the acquisition of this central PP tripotent configuration and to further enrich differentiation into ductal lineage using ductal growth media^33^. Ductal cells were of interest to us in terms of high abundance of CFTR protein in ductal epithelial cells^34^, involvement of ductal pathology, and duct-restricted *CFTR* variants in the development of RAP and CP. The iPSC-based modeling helped us overcome the limitation of obtaining very rare human pancreatitis samples, while capturing the driver genetic composition of pancreatitis patient. This model allowed us to test the specifics of ductal pathology one of which we hypothesized to be the presence of an aberrant hyper mucinous state in ductal cells based on the presence of a dominant mucus phenotype in CFTR-associated genetic disease cystic fibrosis (CF)^35^ (**Figure 1a**). In this study, we compared the isolation and characteristics of primary and iPSC-derived pancreatic ductal cells from a 35-year-old female patient with idiopathic RAP (PANx line) and an age-and gender-matched control (Con). The PANx line had the following genetic mutations: In the 12 gene panel genetic testing for pancreatitis, this patient was heterozygous for a common variant of a large *CFTR* haplotype c.1408G>A (p.Val470Met, rs213950) and a silent mutation c.2562T>G (p.Thr854=, rs1042077) in *CFTR* likely in cis with both variants increasing the odds for pancreatitis when in combination with other disease causing variants. The *CFTR* variant V470M is polymorphic at the amino acid level, and variants at this locus can reduce CFTR activity by up to 60–80%^36^. This variant is also highly prevalent among general population as well as pancreatitis patients with no pathologic association in isolation and can be identified in 40–60% of individuals of all ethnicities^37^. The patient was also heterozygous for the allelic variant ‘T’ at rs7512462 in the gene *SLC26A9*, an important modifier of CFTR function. Notably, this patient was homozygous for the risk factor variant in *APOB*, c.13013G>A (p.Ser4338Asn, rs1042034). This common variant is associated with increased serum triglyceride levels that does not cause pancreatitis in isolation but increases the risk of pancreatitis.

**Figure 1.**
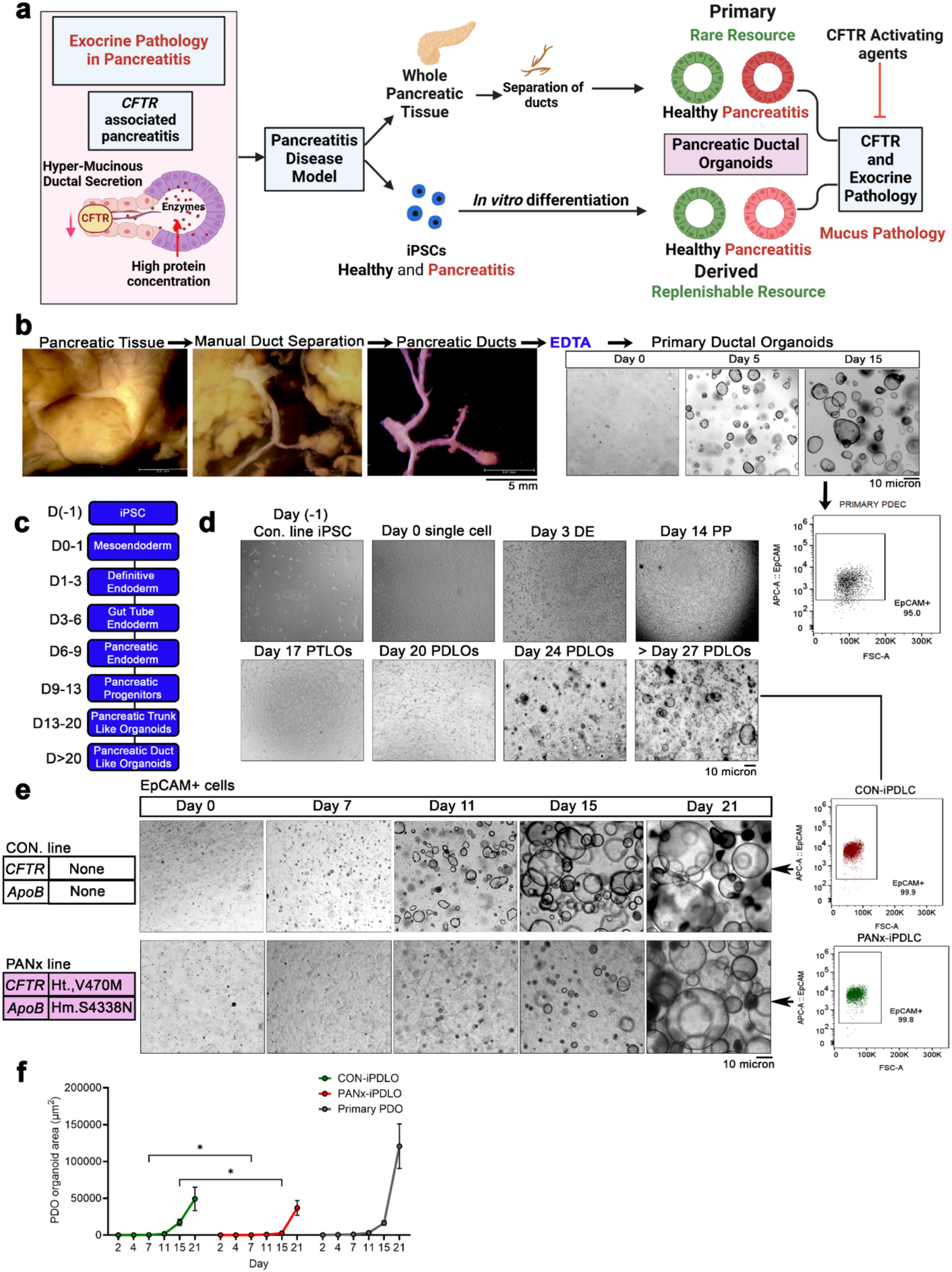
Pancreatitis model of human ductal tissue containing a common *CFTR* variant. **a.** Diagrammatic representation of the events hypothesized to be related to deficit in CFTR function due to the presence of CFTR variants in idiopathic pancreatitis. A lack of CFTR function leads to insufficient flushing of the acini compartment causing accumulation of digestive enzymes that over time lead to acinar and duct damage i.e. autoproteolysis, progressive inflammation and pancreatitis. To investigate CFTR-related pathology in pancreatitis, human model systems, which can be sourced from primary tissues i.e., pancreatic resections are required. Alternatively, the differentiation of iPSCs into pancreatic structures such as ductal tissues is a highly feasible and readily available approach to study ductal pathology including mucus related defects in pancreatitis. **b.** Isolation of ducts from whole pancreatic tissue (non-pancreatitis) followed by EDTA treatment to release stem cells (Day 0) to form ductal organoids that grow and show typical ring-like morphology with large luminal area (Day 15). **c.** Stepwise schemata of the differentiation of iPSCs into ductal progenitors and the formation of pancreatic duct like organoids. **d.** Bright-field images show different stages of iPSC differentiation of a control line and a patient line, PANx, with idiopathic pancreatitis and a common *CFTR* variant into pancreatic duct like organoids (iPDLO). Upon single cell isolation and characterization using flow-cytomtery, these cells were purely epithelial (i.e. ∼99%) based on the presence of the canonical epithelial marker Epithelial cell adhesion molecule (EPCAM). The single cells could be further seeded to generate ring-like epithelial structures i.e. pancreatic ductal like organoids. **e.** Growth characteristics i.e., total area of primary and iPSC-derived pancreatic ductal like organoids (Control and PANx) over extended periods of culture.

To isolate primary ductal organoids (PDO), we either separated ducts from a whole pancreatic tissue if available or ductal fragments from pancreatic cell remnant from TPIAT samples as described earlier^18^. In **Figure 1b**, we isolated ductal tube from whole pancreatic tissue by performing manual microdissection under the stereoscope, followed by EDTA treatment to release stem cells for organoid growth in Matrigel to generate pancreatic ductal epithelial cells (PDEC). Robust ring-like morphology could be seen that is typical of ductal organoids^18,32,38^. For iPSC differentiation, we largely followed the protocol reported by Kleger’s group^33^ until the stage of Pancreatic Trunk like Organoid (PTLO) to Pancreatic Duct like Organoid (iPDLO) differentiation where we optimized a different media composition (detailed media composition and reagent concentrations are described in the methods section) which yielded superior quality of ductal organoids (**Figure 1c**). iPSC-differentiated cells were not characterized during the stages of differentiation using any method other than being identified by the typical morphology of ductal rings (**Figure 1d and 1e**) but at the end of the differentiation process (results described in **Figure 2**). At the end of the iPDLO formation and a subsequent passaging at > day 24, we were able to sort the epithelial cell marker Epithelial Cell Adhesion Molecule (EPCAM^+^) population. Nearly 99% of the cells were EPCAM^+^ in both the control and PANx lines suggesting a pure epithelilal composition of the iPSC-differentiated cells (**Figure 1e**). We also compared the growth dynamics of EPCAM^+^-derived PDO of primary as well as iPSC-origin (**Figure 1f**). Although primary organoids exhibited robust growth in the initial 2-4 days, the growth rate of all the organoid types was on par with each other including for the PANx line over extended days of culture > 7 days. However, the PANx-iPDLOs were consistently smaller in size compared to the controls (**Figure 1f**). We used these organoids to isolate single cells for further molecular and functional characterization.

**Figure 2.**
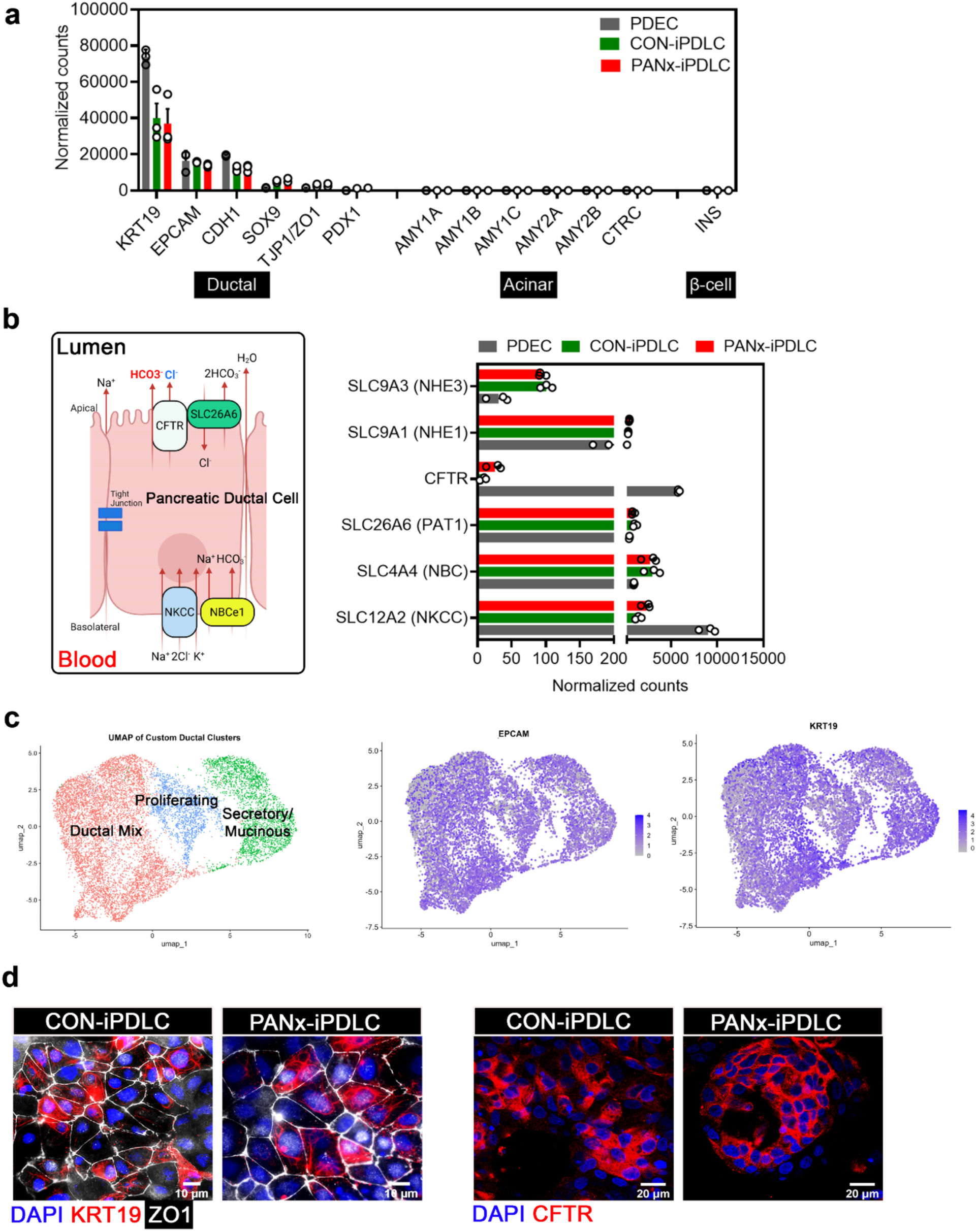
Molecular characterization of iPSC-derived pancreatic ductal tissues. **a.** global transcriptomic data were mined to investigate and present normalized expression counts for pancreatic duct specific markers in iPSC-derived ductal cells from control and PANx line and compared to the primary ductal epithelial cells (PDEC) derived from human ductal organoids. None of the markers corresponding to the acinar cells (*AMY*, amylases) or islets, β-cells (*INS*, insulin) were detected in the primary as well as iPSC-derived ductal tissue and the differentiated structures demonstrated pure molecular characteristics of the ductal tissue marked by *CDH1* (E-Cadherin)*, KRT19* (cytokeratin 19) and *ZO1* (zonula occludens 1). Each dot represents a single well containing the ductal organoids for the specified samples. Error bars represent S.E.M. **b.** Graphical scheme on the left shows the regulation of bicarbonate and fluid secretion through the acitvitites of various transporters and channels in pancreatic duct epithelial cells. The bar-graph on the right depicts mRNA expressions for various transporters including CFTR in iPSC-derived ductal tissues i.e., iPDLC from control and PANx lines and compared with PDEC. Each dot represents a single well containing the ductal organoids from which single cells were prepared for mRNA analysis in the specified samples. Error bars represent S.E.M. *P-value* using one-way ANOVA. **c.** scRNA-seq identifies three major populations of cell clusters: ductal mix (four subclusters), proliferating and secretory/mucinous (two subclusters). scRNA-seq data was analyzed using the Seurat v5 package following the standard pre-processing workflow. Doublets were identified and removed prior to downstream analysis. Quality control (QC) was performed to retain only high-quality cells, excluding those with more than 10% mitochondrial gene expression. The data were then normalized, and highly variable features were identified. Principal component analysis (PCA) was conducted, followed by non-linear dimensional reduction using UMAP to visualize cell clustering and transcriptional heterogeneity. The UMAP projection in scRNA-seq of control and PANx cells vividly illustrates that nearly all cells exhibit positive expression for both KRT19 and EPCAM. This ubiquitous co-expression strongly indicates a predominant pancreatic ductal signature, reflecting a fundamental ductalc identity among the sampled cells. The result underscores the robust ductal lineage characteristics within the analyzed cell population and provides critical insights into pancreatic ductal cell’s composition. **d.** Immunostaining of the ductal epithelium markers cytokeratin 19 and ZO1 and CFTR in control and PANx iPDLC. DAPI was used as a counterstain.

### iPSC-derived control and PANx cells expressed all key pancreatic ductal cell markers

Here, we first characterized the molecular signature of control and PANx ductal cells at > Day 27 of iPDLO formation and post-passage two. Based on global-transcriptome sequencing, primary as well as iPSC-derived ductal cells showed high expression of cytokeratin 19 (*KRT19*), E-cadherin (*CDH1*), zonula occludens 1 (*ZO1*) or tight junction protein 1(*TJP1*), and adult stem cell marker transcription factor SRY-Box Transcription Factor 9 (*SOX9*) and low expression of β-cell enriched and ductal cell pancreatic and duodenal homeobox 1 (*PDX1*). No marker gene expressions for acinar cells (amylases, *AMY*) and β-cells (insulin, *INS*) were present in the differentiated cells from both lines (**Figure 2a**). These expression patterns were very similar to those of the primary ductal organoids and there was no difference in the expression of these markers between the control and PANx cells (**Figure 2a**). The expression of *CFTR* was also confirmed along with other key transporters for bicarbonate secretion in the control and PANx iPDLCs albeit at a lower level for *CFTR* compared to the primary cells (**Figure 2b**). Using single-cell sequencing, we confirmed that iPSC-derived differentiated cells were homogenously ductal (KRT19^+^, EPCAM^+^) and formed three major clusters with seven subclusters that we named ductal mix (composed of mature ductal (cadherins rich), likely endocrine adjunct or transitioning to endocrine based on several overlapping genes expressed in β-cells, and ductal cells undergoing nuclear transformation/DNA repair), actively proliferating and of secretory/mucinous phenotypes (**Figure 2c, Supplementary Figure 1a and 1b**). The presence of ductal markers corresponding to ZO1, KRT19 and CFTR was confirmed at the protein level using immunofluorescence with no detectable difference of expression between the control and PANx cells (**Figure 2d**).

### Pancreatitis associated *CFTR* variant caused reduced CFTR activity that could be restored using CFTR-activating agents

Given that our goal is to understand the ductal pathology due to CFTR-related defect as anticipated for the *CFTR* variant in the PANx line, it become impertive to evaluate the function of CFTR in the PANx cells and compare it with control cells. It was also crucial to conduct this experiment to ensure the formation of a functional ductal epithelium from iPSC-differentiation. CFTR function was evaluated using whole-cell patch clamping in single primary, control, and PANx cells. Primary cells tended to have higher CFTR function than control iPDLCs, but the difference was not statistically significant over the number of independent experiments that were performed (**Figure 3a and 3b**). None of the single patch-clamped PANx cells demonstrated CFTR function corresponding to a total current density of < 0.15 pA/pF and no detectable CFTR-inhibitor-172 (CFTRinh-172) sensitive response while 70% of the control cells showed clear CFTRinh-172 sensitive response upon stimulation using cyclic AMP (cAMP) or cAMP agonist forskolin (FSK) with a median current density of 2.68 pA/pF (**Figure 3c-3e**). We further explored the possibility that the functional consequences of the mutation may be treatable with *CFTR* targeted therapies. Ivacaftor or VX-770 (Brand name: Kalydeco®) is a clinically approved drug used to treat CF in individuals with specific mutations involving gating and conductance defects in the CFTR protein^26^. Ivacaftor is a potentiator that directly binds to CFTR and helps open the channel^39,40^. The binding site is now known based on the cryo-EM structure of ivacaftor in complex with phosphorylated CFTR^41^. Ivacaftor docks into a cleft formed by transmembrane (TM) helices 4, 5, and 8. The predictable binding site is at the signature hinge region in TM8 of the CFTR. The extracellular region of TM 8 rotates around this hinge upon ATP binding, which can be stabilized by ivacaftor. Ivacaftor is known to open even normal CFTR channels and can restore sub-optimal CFTR function as well. Therefore, it can be evaluated as a CFTR-activating agent without the disease pathology for CF and its benefit has been established in a few CFTR-related non-CF conditions^39,40^. Unlike other CFTR activating agents, ivacaftor is a specific potentiator of CFTR activity and hence more suitable agent for evaluating the benefits of CFTR activation and CFTR-directed therapies in human diseases. Also, since PANx cells expressed CFTR protein but showed a deficit in function, this line is suitable to test ivacaftor. Consistent with our expectation, PANx cells exhibited a remarkable rescue of CFTR response to VX-770 treatment in 78% of the tested cells with a median response of 2.97 pA/pF (**Figure 3c-3e**). The functional heterogeneity of the cells for CFTR is expected in asynchronous non-clonal single cell preparations, meaning that not all cells will have functional CFTR leading to a high degree of variability in this assay. To overcome this limitation, we sorted to measure macroscopic CFTR-inhibitor responsive short-circuit currents. These currents were stimulated using FSK stimulation and calculated as *ΔIsc* for iPDLC that were cultured as monolayers on semipermeable transwell supports and formed highly resistant epithelium within 7 days of culture (1500-2000 Ω/cm^2^). We observed similar CFTR-mediated *Isc* in primary and control ductal cells at ΔIsc ± SD of 4.76 ±1.32 and 4.22 ± 0.34 µA/cm^2^, respectively, while PANx demonstrated significantly low but positive CFTR ΔIsc ± SD 1.33 ± 0.75 µA/cm^2^ (**Figure 3f and 3g**). Consistent with earlier results, VX-770 significantly improved CFTR ΔIsc in PANx cells to 2.6 ± 0.9 µA/cm^2^. Another highly adopted method to measure CFTR-dependent function is monitoring FSK-elicited fluid secretion or FSK-induced swelling (FIS) in the 3D Matrigel-embedded organoid system calculated as total organoid area post-FSK stimulation^42,43^. Using FIS, we were able to recapitulate the reduced function of CFTR in PANx iPDLO with an estimated 50% reduction in function and a rescue of function using VX-770 (**Figure 3h and 3i**).

**Figure 3.**
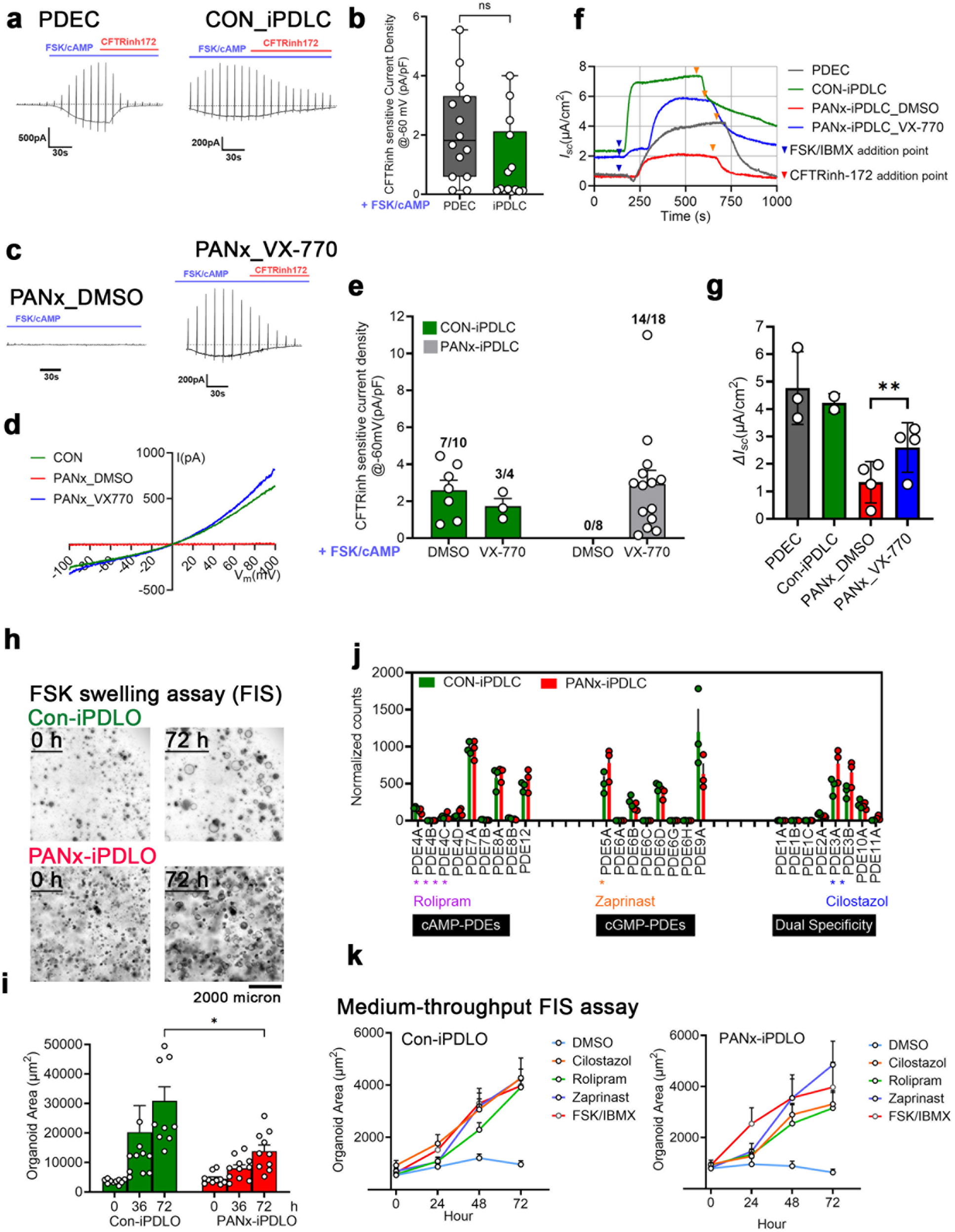
Pancreatitis associated *CFTR* variant caused reduced CFTR activity that could be restored using CFTR-activating agents. **a.** Whole-cell recordings from primary PDEC and control iPDLC demonstrating CFTR-inhibitor sensitive currents elicited using FSK (2 µM). **b.** Bar-graph representing iPDLC CFTR current density recorded at −60mV in single cells isolated from primary PDEC and control iPDLC obtained from whole-cell clamp recording as represented in panel E. The measured whole-cell currents were normalized to cell capacitance. Error bars represent S.E.M. Statistical significance was calculated using student’s t-test. Each dot represents single cell measurement repeated across three to five independent experiments. Error bars represent S.E.M. *P-value* using student’s t-test. ns stands for non-signficant. **c.** Whole-cell recording from PANx iPDLC demonstrating absence of CFTR-inhibitor sensitive currents in response to FSK which were stimulated in the presence of VX-770 (5 µM). **d.** The I-V relationships for CFTR currents in control and PANx iPDLC evoked by 2 µM FSK with or without VX-770 for PANx iPDLC. **e.** Summary of current densities recorded at −60mV for CFTR in control and PANx iPDLCs treated with and without VX-770 in the patch-pipette. The measured whole-cell currents were normalized to cell capacitance. Each dot represents single cell measurement repeated across three to five independent experiments. Error bars represent S.E.M. Statistical significance was calculated using student’s t-test. **f.** Short-circuit current (*Isc*) measurements to measure CFTR function in control and and PANx iPDLCs that were differentiated on transwell cultures and stimulated using FSK (2 µM) or VX-770 in the presence or absence of VX-770 **g.** Bar-graph shows quantitation of the maximal Δ*Isc* n control and and PANx iPDLCs transwell cultures under different treatment conditions. Each dot represents a single experiment. Error bars represent S.E.M. Statistical significance was calculated using one-way ANOVA. **h.** Bright-field images showing fluid secretion in response to cAMP agonists FSK/IBMX called FSK-induced swelling (FIS) in control and PANx iPDLOs at baseline and 72 h post-stimulation. **i.** The bar-graph represents quantitation of fluid secretion based on total organoid area (µm^2^) in the two samples at 0 h and 72 h post-FSK/IBMX treatment. Each dot represents a single organoid, and measurements were repeated across three independent experiments. Error bars represent S.E.M. *P-value* using one-way ANOVA. **j.** Global transcriptomic data were mined to investigate the expression level of various phosphodiesterases (PDEs) in iPSC-derived ductal organoids from control and PANx line. The PDEs can be categorized as cAMP or cGMP-specific or PDEs with dual specificity. PDE inhibitors Rolipram, Zaprinast, and Cilostazol are shown with asterisks to represent their respective specificities towards different PDEs. Each dot represents a single well containing the ductal organoids in the specified samples. Error bars represent S.E.M. **k.** Medium throughput assay with line-graphs representing FIS in control iPDLOs (TOP) and PANx iPDLOs (BOTTOM) in response to simultaneous treatment with PDE inhibitors rolipram (20 µM), zaprinast (50 µM) and cilostazol (20 µM). No FSK treatment with DMSO alone was used as no swelling response control. Error bars represent S.E.M.

Given that CFTR is a cAMP-regulated chloride channel, there are additional FDA-approved drugs that are in clinical use for various human conditions that alter cAMP signaling, including inhibitors of the cAMP and cGMP degrading enzymes-phosphodiesterases (PDEs). Based on transcriptomic analysis, cAMP-degrading PDEs, such as PDE4A, PDE7A, PDE8A and PDE12, cGMP-degrading PDEs PDE5A, PDE6B, PDE6D and PDE9A, and dual-specificity PDEs PDE3A, PDE3B and PDE10A were abundantly expressed in control and PANx iPDLCs (**Figure 3j**). The most notable finding is the relatively high abundance of cGMP degrading enzymes-PDE5A and PDE9A. Rolipram is a selective PDE4 inhibitor that has been researched for its benefit in the treatment of Alzheimer’s Disease and inflammatory respiratory conditions such as asthma^44^. Zaprinast is a precursor to the drug sildenafil (Viagra), which is a PDE-5,6,9 and 11 inhibitor and is in clinical use for the treatment of erectile dysfunction^45^. It was originally considered as a treatment for allergic asthma and cardiovascular diseases^46,47^. Cilostazol is a potent PDE3A inhibitor that is used to treat periphery artery disease via vasodilation through cAMP generation^48^. Given the abundance of select cAMP and cGMP PDEs in iPDLC, which are targets for the above-described drugs, we tested the effect of these compounds on CFTR function in the control and PANx iPDLO using FIS assay as this assay is amenable to medium throughput testing for multiple drugs. Zaprinast and cilostazol in combination with FSK generated most robust CFTR FIS response in PANx iPDLO that did not plateau after 72 h of measuring fluid secretion response vs. FSK alone (**Figure 3k**). The control iPDLO also responded positively to zaprinast and cilostazol treatment relative to the FSK treatment alone (**Figure 3k**). These data suggest that PDE inhibitors can be considered as therapeutic candidates for restoring CFTR functional defects during pancreatitis.

### Ductal cells with pancreatitis exhibited a hypermucinous state and other pathological features that were reversible in the presence of ivacaftor treatment

Given that CFTR activity is directly related to the mucus phenotype in various CF organs^49^, we investigated the possibility of mucus pathology in PANx cells. Based on the global transcriptomic analysis, there was a multi-fold increase in mucin levels, with a significant increase in membrane-bound mucin 1 (*MUC1*) and secreted mucin 5AC (*MUC5AC*) in PANx cells compared to the control cells (**Figure 4a**). Only majorly expressed MUCs in ductal cells were plotted. One prominent example of the mucin biosynthesis pathway is Trefoil factor 1 (*TFF1*), which is synthesized by the surface mucous cells across the gastric epithelia and is known to be co-secreted with MUC5AC to aid in the formation of mucous layer^50,51^. Both *TFF1* and *MUC5AC* were upregulated by more than 10-fold and 200-fold respectively, in PANx cells (**Figure 4a**). Galectin 3 (*LGALS3*) is another gene that was highly upregulated at > 4-fold change in PANx cells (**Figure 4a**) and plays a role in stabilizing the mucus layer by interacting with the sugar moieties on the mucins^52^. Another contributing genetic factor to the mucus phenotype was the highly upregulated expression of a protease inhibitor WAP four-disulfide core domain protein 2 (*WFDC2*) in PANx cells (**Figure 4a**) which contributes to the secretion and formation of mucus barrier and is expressed by mucous cells^53^. A significant increase in the expression of REG4 (32-fold) (**Figure 4a**) also suggested an increase in the secretory cell phenotype ^54^. Our single cell analysis also demonstrated enrichment for mucinous ductal cell population in PANx ductal cells marked by MUC5AC, MUC5B and MUC1 within the mucinous population (**Figure 4b**). At the level of the protein, there were much higher levels of secreted MUC5AC in PANx cells vs. control under reducing conditions using dithiothreitol (**Figure 4c**). Importantly, the expression levels of both the transcripts as well as secreted protein for MUC5AC were restored to the control values upon 24-48 h treatment with VX-770 (**Figure 4a and 4c**). Using a biotinylated mucin binding probe (Mucin Probe, biotin-StcE) and epithelial cell-specific stain detecting CDH1, we identified large cystic structures in PANx transwell differentiated cultures that also showed larger nuclei exhibiting the characteristics of mucinous neoplasms (**Figure 4d-4f, Supplementary Figure 2a**). Remarkably, these structures largely resolved upon treatment with VX-770 (**Figure 4c-4f**). The PANx line exhibited distinct mucous cells and membrane-bound mucins that were in complete absentia in control cells and disappeared upon treatment with VX-770 (**Figure 4d and 4f**). Intraductal mucinous pancreatic neoplasms are mucin-producing cystic lesions of the exocrine pancreas, usually localized within the pancreatic ducts^55,56^. Dysregulated activities of *KRAS*, *CDKN2A*, *TP53*, and *SMAD4* are strongly associated with the development of neoplasms^57,58^. In the PANx line, tumor suppressor *TP53* expression was significantly downregulated in the PANx line and was restored to the control level in the presence of VX-770, whereas *KRAS*, *CDKN2A* and *SMAD4* were not altered (**Supplementary Figure 2c**). Approximately 15% to 30% of ductal neoplasms may be malignant^59^. The primary marker used to identify malignant mucinous pancreatic neoplasms is carcinoembryonic antigen cell adhesion molecule 1/5 (CEACAM1/5) secreted into the ductal fluid^60^. Unfortunately, the patient line demonstrated exceedingly high levels of *CEACAM5* at the mRNA level compared with the control line suggesting likely malignancy which remain clinically undiagnosed in this patient as of now (**Supplementary Figure 2b and 2c**). The *CEACAM5* levels were also restored to the normal levels in the presence of VX-770. DKK1, an antagonist of Wnt/β-catenin signaling, was downregulated by >4-fold in PANx cells, indicating an increased migratory effect and potential role in tissue metastasis ^61^ and VX-770 had a marginal effect on increasing its levels (**Supplementary Figure 2c and Figure 5d**). Therefore, not only the restoration of defective CFTR function, but we also established the multi-layered benefits of VX-770 on pancreatitis that could be due to CFTR-related primary or secondary events.

**Figure 4.**
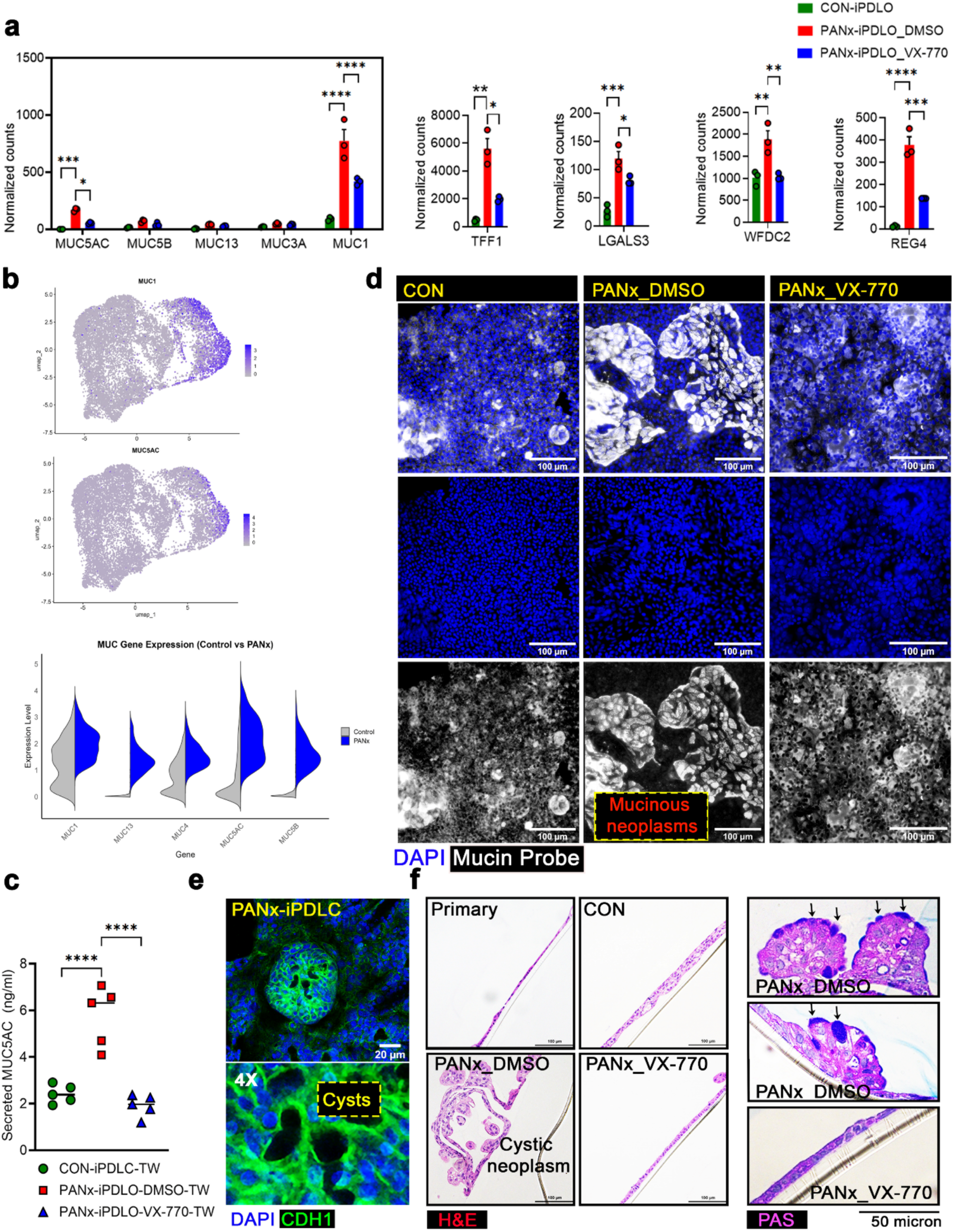
Ivacaftor reversed mucinous pathological feature in pancreatitis ductal cells. **a.** Normalized expression counts obtained from total transcriptomic data for different major mucins as well as for the regulator of mucins including TFF1, WFDC2 and REG4 in control and PANx iPDLC (± VX-770 (5 µM, 24 h)). Each dot represents a single well containing the ductal organoids in the specified samples. Error bars represent S.E.M. *P-value* was calculated using multiple t-test. **b.** The UMAP visualization for sc-RNAseq data in control and PANx cells revealed that the expression of mucin genes was identified in a unique mucinous-ductal cell population. This spatially confined expression pattern supports the identity and functional specialization of this subset, indicating their distinct role in mucin production and stabilization within the ductal cell population. Violin plots comparing control and PANx conditions for sc-RNAseq data revealed elevated expression of major mucin genes within the mucinous ductal cell population in PANx iPDLC alluding to the presence of a hypermucinous state. **c.** MUC5AC-specific ELISA was used to measure the amount of secreted protein into the media collected from the transwell differentiated cultures of control and PANx iPDLC. Each dot represents a single experiment. Error bars represent the S.E.M. Statistical significance was calculated using one-way ANOVA. **d.** Immunostaining for total mucin using a mucin probe (Biotin-STcE) in the transwell differentiated cultures of control and PANx iPDLC. DAPI was used as nuclear counterstain. 3D Mucinous neoplasms were a structural characteristic of differentiated PANx cells, and these structures remarkably disappeared following treatment with VX-770 (5 µM, 24 h). **e.** Confocal images showing positive CDH1 immunostaining for cystic neoplastic structures observed in PANx cells indicating their epithelial origin. **f.** H&E staining shows cystic neoplasms that were only identified in differentiated PANx cells and remarkably disappeared following treatment with VX-770 (5 µM, 24 h). PAS staining shows the presence of distinct mucinous cells and membrane-bound mucins in PANx cells that were not visible following treatment with VX-770 (5 µM, 24 h).

**Figure 5.**
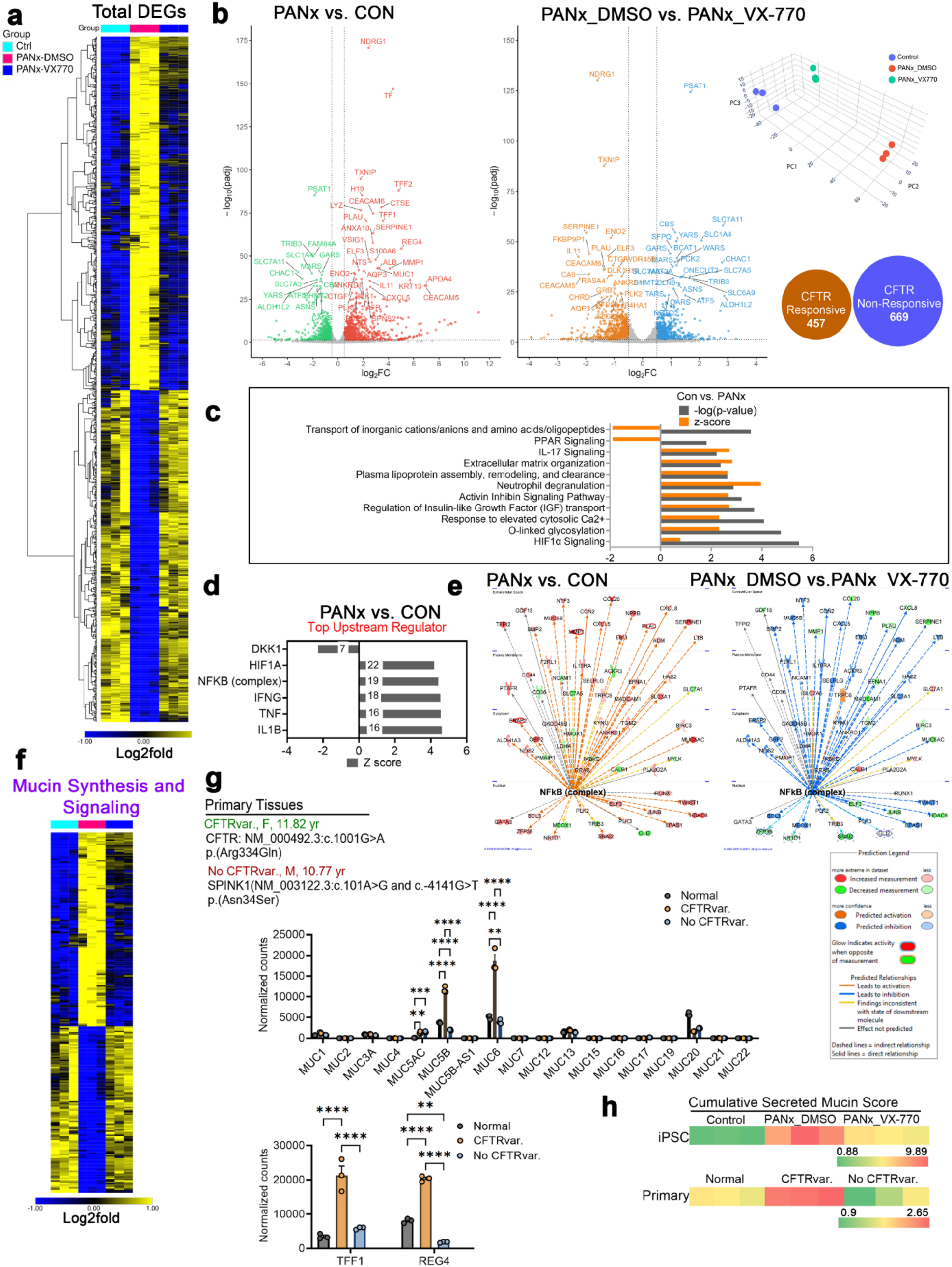
Differential gene expression analysis reveals reversal of molecular signature for ductal tissue pancreatitis using Ivacaftor. **a.** Differential expression gene (DEG) map for three conditions obtained using total transcriptomic data: Control, PANx-DMSO, and PANx-VX-770 showing clustering of grouped samples and distinct expression landscape with a large overlap between the control and PANX-VX-770 sample groups demonstrating reversal of disease signature for PANx using VX-770. Heatmap was generated using Morpheus application (https://software.broadinstitute.org/morpheus/). Further, these genes were seggregated into CFTR-resonsive and non-responsive based on VX-770 reversible signature in PANx cells relative to the control cells. **b.** Volcano-plot showing top 50 DEGs (UP-Right, DOWN-Left) expressed between control and PANx cells as the comparison group of PANx cells either treated with DMSO alone or VX-770 for 24 h. Several top genes showed reversal of expression in the PANx cells upon treatment with VX-770 among which multiple genes belonged to mucins or mucin regulatory genes. Princpal component (PC) analysis plot showing clustering of control and VX-770 treated PANx cells. About 40% of the genes were altered by VX-770 which were differentially expressed in the control cells termed as CFTR responsive with 60% DEGs not affected by VX-770 termed as CFTR non-responsive. **c.** Canonical pathway enrichment revealed top altered pathways (up and down based on z-score) in PANx cells including for O-linked glycosylation which is directly linked to increased synthesis of glycoproteins including mucins and Hif1α signaling directly implicated in pancreatitis assoictaed tissue injury. A highly upregulated pathway signature for neutrophil degranulation suggests ability of PANx cells to promote inflammation. **d.** Top upstream regulators activated or inhibited in PANx cells. Analysis was performed using IPA, Qiagen©. **e.** Activated NFƙb signaling node in PANx cells with a clear reversal of activated signature in VX-770 treated PANx cells. Analysis was performed using IPA, Qiagen©. Identifying legends are included on the bottom right. **f.** DEGs for mucin biosynthesis based on O-linked glycosylation database in lung mucous cell for three conditions obtained using total transcriptomic data: Control, PANx-DMSO, PANx-VX-770 showing enrichment for mucin biosynthesis in PANx that was reversible using VX-770. **g.** Gene expression analysis from whole-transcriptomic sequencing in primary ductal tissues obtained from pediatric patient’s remnant TPIAT samples with and without a *CFTR* variant reproduced hypermucinous and secretory profile in pancreatitis samples with the variant similar to the iPSC-derived tissues. The hypermucinous phenotype is additionally supported by highly upregulated expression of TFF1 and REG4 in the cells with *CFTR* variant vs. no variant. Statistical significance was calculated using one-way ANOVA. **h.** Heat-map shows cumulative normalized count score for secreted MUC genes (sum for MUC2, MUC5AC, MUC5B, MUC6, MUC7 and MUC19 and normalized to the control/normal) across iPSC-derived ductal tissues vs. primary tissues under different conditions.

At the global transcriptomics level, there were a total of 940 differentially expressed genes (DEGs) for PANx: 485 up and 455 downregulated genes (**Figure 5a-5b**). At log2 fold threshold of 1, 41% of the differentially altered genes in PANx vs. control were affected by VX-770 treatment. We labeled these as CFTR activity-responsive genes (**Figure 5b**). Among these, genes associated with mucin biosynthesis (51/940 DEGs; 49/51 up and 2/51 down) and inflammation (107/940 DEGs) as anticipated for the PANx line were significantly affected, and this signature was notably reversible in the presence of VX-770 (**Figure 5f, Supplementary Figure 3a**).

A general view of the pancreatitis pathology is that it is caused by a combination of auto-digestive reactions due to premature activation of trypsinogen within the pancreatic acini and a parallel inflammatory cascade due to the recruitment of macrophages and granulocytes both leading to tissue damage^62,63^. Ductal and acinar cells themselves release several pro-inflammatory cytokines, such as interleukin (IL)-1, IL-6, IL-8, IL-18, IL-33, and tumor necrosis factor (TNF)-α to start a vicious process of pancreatic injury and inflammation and eventually systemic injury that can lead to death^64,65^. Sustained intra-acinar activation of nuclear factor-κB (NF-κB) downstream of initial injury appears to be the triggering event in the development of inflammation in chronic pancreatitis^66^. Indeed, the major pathways that were upregulated in PANx cells included HIF1α and NF-κB signaling which are associated with an increased production of inflammatory cytokines (**Figure 5c and 5d, Supplementary Figure 3a**). Treatment with VX-770 had remarkably reverted this signature in the PANx cells with several genes affected by VX-770 as mapped in the NF-κB signaling network (**Figure 5e**). We also listed the key cytokines that were upregulated in the PANx line (**Table 1**) many of which have already been established as pro-inflammatory and are known to play a role in pancreatitis including IL-11, IL-33 and several members of the C-X-C motif chemokine (CXCL) family (CX3CL1, CXCL1, CXCL12, CXCL16, CXCL5, CXCL8). Among the notable pro-inflammatory cytokines, following cytokines levels were reduced by VX-770: CCL20, CCL28, CXCL8 and IL-11 (**Table 2**). The PANx line showed an abundance for signature corresponding to the formation of pancreatic lesions (260/940 DEGs; 140/260 up and 120/260 down) (**Supplementary Figure 2b**) as was morphologically and immunohistochemically captured in the earlier observations. Due to the presence of APOB mutation in the PANx line, we also examined pathways for lipid homeostasis in the cells (**Supplementary Figure 3b**). S4338N *APOB* mutation is known to be associated with hypercholesterolemia, hypertension, atherosclerosis and coronary heart disease. DGE analysis suggested that a total of 69 genes were altered in the lipid biosythesis pathway (69/940 DEGs) that were affected by VX-770 (**Supplementary Figure 3b**). However, there was no clear indication of abberration of lipid homeostasis in PANx cells based on pathway analysis.

We used primary ductal pancreatitis tissue obtained from TPIAT remnant samples to validate our data from iPSC-derived ductal tissues regarding mucinous pathology and its association with CFTR. A similar hypermucinous signature was present in ductal cells with *CFTR* variants compared to those without variants and only pancreatitis-related *SPINK1* mutation (**Figure 5g**). This signature was characterized by an increased transcriptional profile for several mucin genes, predominantly MUC5B, and an enhanced secretory cell program regulated by TFF1 and REG4. The secreted mucin score was calculated for primary and iPSC-derived ductal tissues based on cumulative counts of secreted mucins (MUC2, MUC5AC, MUC5B, MUC6, MUC7, and MUC19), showing an overlapping trend of enhanced mucin secretion with the iPSC-derived PANx ductal cells and specific to the state of presence ot absence of a *CFTR* variant (**Figure 5h**). Therefore, this data strongly supports the presence of a common hypermucinous state across CFTR-related pancreatitis and demonstrates the robustness of our iPSC model system in recapitulating this pathology.

### Modeling hyperamylasemia as a function of hyperviscous ductal fluid using a microsystem device

The loss of acinar function is a known feature of pancreatitis and could be a consequence of acinar damage^67,68^. Hyperamylasemia, which is defined as elevated serum amylase levels >110U/ml, is a common finding in up to 75% of cases of pancreatitis^69^. Hyperamylasemia is primarily caused by damaged acini in pancreatitis^70^. Elevated serum amylase is routinely used as a biomarker for acute pancreatitis attack. Elevated amylase levels have earlier been reproted to be consistently associated with the presence of *CFTR* and *SPINK1* mutations^71^. There is also a direct correlation between elevated amylase and low pH of the pancreatic juice^72^. Periodic Acid-Schiff (PAS) staining in pancreatic tissues revealed a high secretory state in ductal epithelial cells, characterized by glycoproteins, including mucins (**Figure 6a**). These glycoproteins contribute to the composition of ductal secretions, which are released through a branching network of ducts originating from the main duct. This network forms interlobular and intralobular duct systems that extend to the terminal acinar compartments. We hypothesized that more viscous ductal fluid composition due to a deficit in CFTR function causes insufficient flushing of acini and accumulation of digestive enzymes, leading to acinar damage and thus increased amylase accumulation or release. To artificially enhance the viscosity of ductal fluid, we used 10% glycerol which is known to have no apparent impact on cell viability by itself and be often used as a cryopreserving agent. We utilized a commercially available chip from Ibidi that enabled communication between two chambers via a narrow channel (**Figure 6b**). We seeded primary acini freshly isolated from pancreatic tissues with no apparent defects in isolated acinar activity in one chamber and applied 60 µl of the apical fluid from control ductal cells differentiated on TW as is or mixed with 10% glycerol while maintaining the effective composition of ductal fluid at 90% under both conditions. After 24 h of incubation, we performed amylase activity assay to assess acinar cell function. Acinar cells exposed to a more viscous fluid secreted ∼1.5-fold higher amylase compared to the ductal fluid without glycerol (**Figure 6c**). We next assessed whether increased viscosity was associated with acinar cell damage. Cells exposed to viscous ductal fluid exhibited ∼3-fold higher LDH activity, suggesting damage (**Figure 6d**). Since glycerol can act as a weak base, we checked media pH at the concetration of 10%, glycerol and it did not cause any pH change in the media. These data suggest that fluid viscosity has an impact on acinar cell condition, and this is a likely pathological scenario in the case of reduced CFTR activity and related increase in ductal fluid viscosity during the development of pancreatitis.

**Figure 6.**
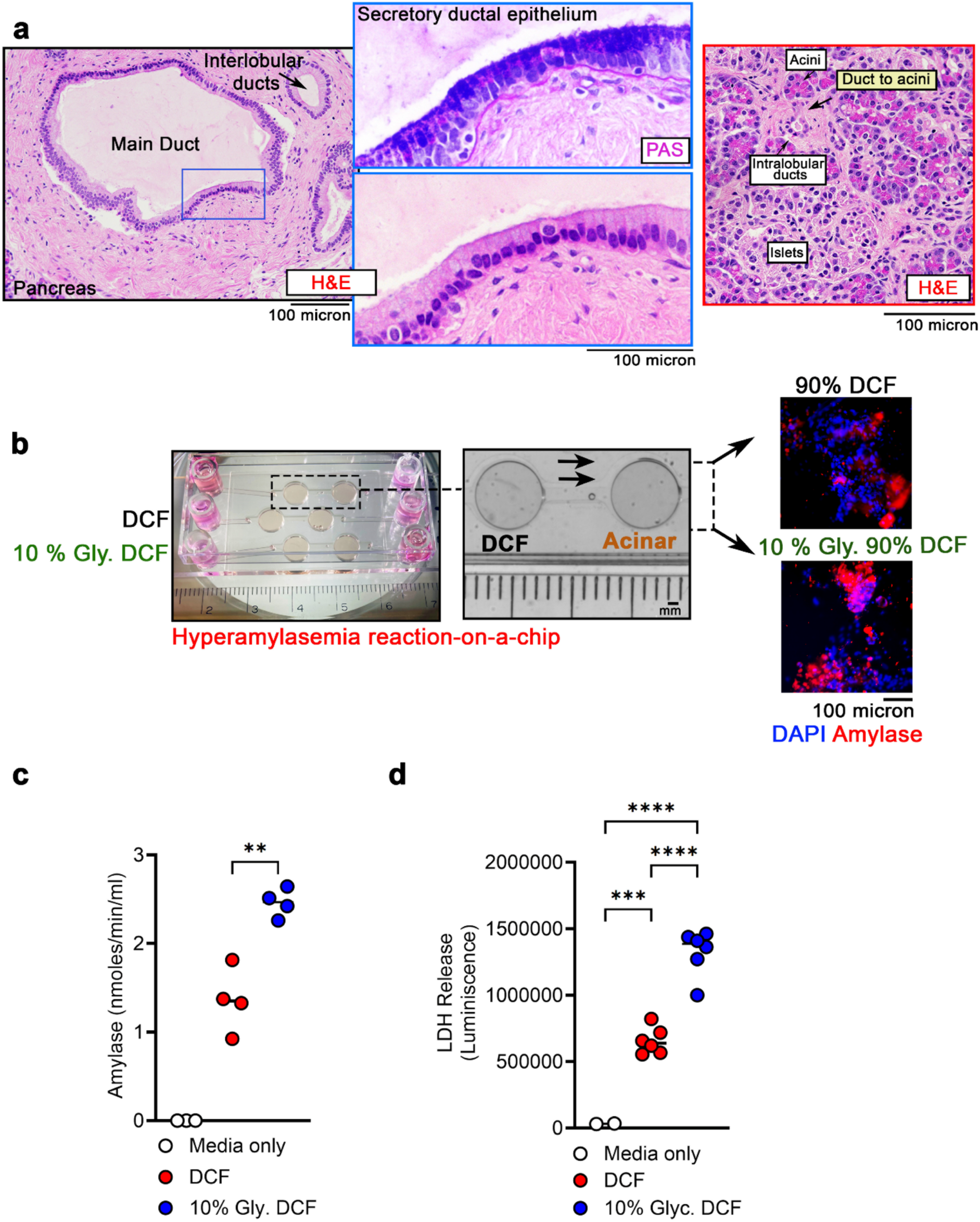
Modeling the reaction for hyperamylasemia using a microsystem device. **a.** H&E images of pancreatic tissue showing main duct and interlobular ducts. The pancreatic duct epithelium is highly secretory marked by glycoprotein rich cells which were detected using PAS stain. The intralobular ducts connect to the acini and regulate their function by delivering protein rich ductal secretions. **b.** A microsystem device from Ibidi containing two chambers connected by a narrow channel was used to investigate the effect of the viscous ductal cell fluid (DCF) on acinar cell function. Chamber on the right was used to culture primary acinar channels, and ductal fluid was collected from normal control iPDLC differentiated on transwells. To increase viscosity, the ductal fluid was mixed with glycerol at 10% concentration while keeping the effective ductal fluid composition at 90% same in the samples with and without glycerol. Amylase specific immunistaining on the right shows acinar cells seeded on the microsystem device under the treatment conditions of 100% DCF and 90% DCF,10% glycerol. **c.** The bar-graph represents amylase activity in the fluid collected from the acinar chamber in the samples exposed to complete ductal medium and ductal cell fluid containing 10% glycerol compared to media only. Each dot represents a single sample averaged from a duplicate across independently performed experiments. Error bars represent the S.E.M. Statistical significance was calculated using student’s t-test. **d.** The bar-graph represents lactate dehydrogenase (LDH) activity in the fluid collected from the acinar chamber to assess acinar cell condition in the samples exposed to complete ductal medium and ductal fluid containing 10% glycerol compared to media only. Each dot represents a single sample averaged from a duplicate across independently performed experiments. Error bars represent S.E.M. Statistical significance was calculated using student’s t-test.

## Discussion

Pancreatitis is a debilitating condition that significantly impacts quality of life and progresses rapidly due to poor prognosis and limited strategies for pain management. Pancreatitis-related studies are limited because of the lack of tangible model systems that recapitulate key pathological features of pancreatitis. Using iPSC-derived modeling, we were able to recapitulate clear pancreatitis-related pathology in the ductal epithelium derived from a patient with idiopathic pancreatitis who carried a common pancreatitis-related *CFTR* variant. Importantly and surprisingly, the pancreatitis signature is inherently present in these cells without an overt injury. Nearly 40% of patients with pancreatitis are known to carry various *CFTR* variants. Several hurdles have been identified in the establishment of an early diagnosis as well as clear pathological course for pancreatitis. First, there are several non-specific symptoms of pancreatitis that overlap with other conditions^73,74^. Second, diagnostic imaging of the pancreas remains inconclusive in many cases and is highly variable^75^. Third, specific biomarkers for diagnosing pancreatitis in early stages of development are lacking^76^. Fourth, several genetic alterations or environmental factors may be involved, leading to complex pathology, and multiple tissue systems are usually impacted in the disease development. The development of human model system remains central to understand the aspects of early pathogenesis of the human disease and cell-type specific pathology, and to test and develop therapies. By generating pure pancreatic cell-types, more complex multi-tissue systems can be evolved by precise cell-mixing to understand the involvement of multiple cell-types and specific cell-cell communication in disease development. We established that using an iPSC-derived model system of pancreatic ductal cells, deficit in CFTR function within the ducts can drive pathology of idiopathic pancreatitis in the patients with a common CFTR variant. This is a contrasting finding for the commonly held notion for acini as the site for initial pancreatitis injury^63^. This finding is further supported by the data that by rescuing CFTR function using FDA approved CFTR modulator ivacaftor several pancreatitis associated pathological characteristics that align with the clinical features of pancreatitis could be reversed. Nearly, 40% of the pancreatitis-related ductal cell genes responded to ivacaftor treatment suggesting the presence of a major CFTR associated pancreatitis signature.

In CF, CFTR protein product is defective and associated with devastating lung infections and other extra-pulmonary manifestations including pancreatic insufficiency and meconium ileus^35^. A major pathological feature of CF in these organs is the development of mucus plugs and hyperviscous conditions in the airway, pancreas, and gut^35,77^. A deficiency in fluid secretion alters the mucus properties resulting in the failure of attachment from the mucous cells^78^. Viscosity is an important parameter for fluid that is secreted into the tubular configurations of the airway, pancreas, and gut for lubrication and, flow, flushing, dilution, and release of the luminal factors. Fluid properties are significantly altered in CF and that was the hypothesis that we tested in this pancreatitis model due to CFTR-related defect. To our surprise, the mucus defect was predominantly associated with the development of pancreatic neoplasms that were hypermucinous and developed in the transwell differentiated cultures of the pancreatitis line. This pathological transformation is the common sequale of chronic pancreatitis. This mucinous pathology is distinct from CF which is normally characterized by goblet cell hyperplasia and abnormal sticky mucus^79^. We exposed normal acinar cells to a hyperviscous environment using glycerol on a chamber device composed of two compartment interactions through a narrow channel, we were able to model hyperamylasemia characterized by upregulated production of amylase associated with acinar cell damage. The viscosity was simulated using 10% glycerol, which is known to have no effect on cell viability. The acinar cells maintained in both environments i.e. normal ductal fluid and ductal fluid plus glycerol did not show any apparent difference in cell quality and morphology. Further, the pancreatitis cells served as the primary source for the expression of CEACAMs that are used as the biomarker for pancreatic cancers^80^ and this feature was reversed by ivacaftor. The role of CEACAMs is suggested to be key in driving the inflammatory response during pancreatitis^81^.

Notable inflammation driving pathways that were significantly high in the PANx cells were NF-κB and HIF-1 signaling^82^ that are known to directly increase the production of various pro-inflammatory cytokines such as TNF-α, IL1β and chemokines during pancreatitis and are also known to be highly elevated in animal models of pancreatitis^83^. The HIF-1 pathway also contributes to the tissue necrosis leading to pancreatic injury. Ductal cell-derived cytokines such as IL-11, CCL20 and CXCL8 were highly upregulated in PANx cells and reverted to normal control levels using ivacaftor (Table 2). These cytokines are known to propagate immune cell infilteration and fibrosis during pancreatitis^84^.

The question arises whether this study is useful for pancreatitis in patients with other mutation types. This study highlights ductal tissue pancreatitis which is foundational for studying pancreatitis in patients with various CFTR variants provided with an evidence of CFTR dysfunction irrespective of the nature of mutation. We found that CFTR functional deficit is associated with key pathological features of pancreatitis. The molecular and morphological features observed with the PANx line overlap with clinically reported features in the general population however, we revealed new layers of information such as drug-reversible disease features in pancreatitis and that CFTR dysfunction in ductal tissue can drive mucinous pancreatic neoplasms, inflammation, and could be associated with pancreatitis attacks. We were able to validate a similar hypermucinous states in primary pancreatitis ductal tissue with a *CFTR* variant which was absent in another paitent sample with no *CFTR* variant and mutation in PRSS1 gene. Therefore, we expect a broad application of these findings to other pancreatitis patients even with differenet set of mutations.

As a model system, iPDLCs exhibited long-term passaging capabilities (15-20 passages) and retained their cellular, molecular, and functional characteristics, unlike primary cells, which are challenging for long-term growth beyond passage 5^18,19^. iPDLCs were suitable for multiple applications, including physiologic assays and expression-based analysis. This study establishes a pipeline for utilizing such a model system for drug development, repurposing, and therapeutic testing. There are major species-driven differences in pancreatic structure and diseases^20,85^. Many therapeutic agents beneficial in mouse models of pancreatitis have not been clinically translated^86^. Using the human-based model system, we established that CFTR-specific agent ivacaftor can reverse several pathological features of pancreatitis, including downstream sequelae such as cancer. Pancreatic cancers consittute the largest risk for death among pancreatitis patients. This approach could be a breakthrough treatment for patients with pancreatitis as we identified hypermucinous neoplasms as reversible pathology in CFTR-related pancreatitis, potentially a primary or secondary event to CFTR functional activation.

### Study approval

This study was approved by the Cedars Sinai Medical Center Institutional Review Board study titled CFTR functional studies, STUDY00001735.

## Acknowledgments

This work was funded by Cedars Sinai Pilot Feasibility award to KA, Cystic fibrosis Foundation, and National Institutes of Health (NIH) awards DK080834 and P30-DK117467 to APN. We thank Dr. Maisam A. Abu-El-Haija, MD, Medical Director, Pancreas Care Center, Cincinnati Children’s Hospital for providing the TPIAT samples. We thank Dr. Dhruv Sareen, Executive Director, Cedars-Sinai Biomanufacturing Center, for help generating control and patient’s iPSC lines. The authors thank Ms. Bonnie Paul for assisting with coordinating patient studies following rigorous regulatory measures for human subject-based studies.

## Footnotes

### Conflict of interest

The authors declare that no conflict of interest exists pertinent to this manuscript.

Methods and supplementary figures and legends are included in the supplementary information.

## Author contributions

The study was conceived by PE, APN and KA and supervised by APN and KA. JL conducted whole cell patch clamp experiments. PL, HL, NT and AGJ analyzed transcriptomic data. DH assisted with the analysis of mucinous pancreatic neoplasms. AF, JDN, BA and SRP reviewed and edited the manuscript and provided key clinical insights into the study. PE and KA wrote the first and several other draft manuscripts before sending a version to all authors to comment on. All authors received, commented on, and approved the submission of this manuscript.

## Tables

Table 1 and Table 2

