## Supplementary Table for "Differential Expression and Microsystem Physiology Reveal Predominant and Drug Reversible CFTR-Related Defects in Idiopathic Pancreatitis"

| Expr Log Ratio | Expr p-value | Symbol | Entrez Gene Name | Location | Type(s) | Drug(s) |
|---|---|---|---|---|---|---|
| -0.629 | 0.00322 | BMP8A | bone morphogenetic protein 8a | Extracellular Space | cytokine | |
| 1.509 | 0.0186 | CCL15 | C-C motif chemokine ligand 15 | Extracellular Space | cytokine | |
| 4.685 | 0.000952 | CCL20 | C-C motif chemokine ligand 20 | Extracellular Space | cytokine | |
| 0.863 | 0.00025 | CSF1 | colony stimulating factor 1 | Extracellular Space | cytokine | PD-360324, lacnotuzumab |
| 0.51 | 0.000853 | CX3CL1 | C-X3-C motif chemokine ligand 1 | Extracellular Space | cytokine | |
| 0.521 | 0.0477 | CXCL1 | C-X-C motif chemokine ligand 1 | Extracellular Space | cytokine | |
| -0.85 | 0.00151 | CXCL12 | C-X-C motif chemokine ligand 12 | Extracellular Space | cytokine | NOX-A12, necuparanib |
| 0.629 | 0.00674 | CXCL16 | C-X-C motif chemokine ligand 16 | Extracellular Space | cytokine | |
| 2.537 | 0.0000175 | CXCL5 | C-X-C motif chemokine ligand 5 | Extracellular Space | cytokine | |
| 1.657 | 0.0000448 | CXCL8 | C-X-C motif chemokine ligand 8 | Extracellular Space | cytokine | BMS-986253 |
| 2.658 | 0.00091 | EBI3 | Epstein-Barr virus induced 3 | Extracellular Space | cytokine | |
| 0.762 | 0.00434 | EDA | ectodysplasin A | Plasma Membrane | cytokine | |
| 2.502 | 0.0000809 | IL11 | interleukin 11 | Extracellular Space | cytokine | 9MW3811 |
| 0.95 | 0.0322 | IL32 | interleukin 32 | Extracellular Space | cytokine | |
| 3.547 | 0.00259 | IL33 | interleukin 33 | Extracellular Space | cytokine | |
| 2.137 | 0.00256 | IL36G | interleukin 36 gamma | Extracellular Space | cytokine | |
| 0.916 | 0.000438 | LIF | LIF interleukin 6 family cytokine | Extracellular Space | cytokine | |
| 1.318 | 0.0366 | LTB | lymphotoxin beta | Extracellular Space | cytokine | |
| 0.898 | 0.00433 | TIMP1 | TIMP metallopeptidase inhibitor 1 | Extracellular Space | cytokine | |
| 1.519 | 0.0018 | TNFSF13B | TNF superfamily member 13b | Extracellular Space | cytokine | belimumab, tabalumab, telitacicept |
| 0.717 | 0.0144 | TNFSF9 | TNF superfamily member 9 | Plasma Membrane | cytokine | |

**Table 1. Cytokines elevated in PANx cells relative to the control ductal cells**

| Expr Log Ratio | Expr p-value | Symbol | Entrez Gene Name | Location | Type(s) | Drug(s) |
|---|---|---|---|---|---|---|
| -1.766 | 0.00293 | CCL20* | C-C motif chemokine ligand 20 | Extracellular Space | cytokine | |
| -1.601 | 0.0425 | CCL28 | C-C motif chemokine ligand 28 | Extracellular Space | cytokine | |
| -1.074 | 0.000712 | CXCL8* | C-X-C motif chemokine ligand 8 | Extracellular Space | cytokine | BMS-986253 |
| -2.211 | 0.0000892 | IL11* | interleukin 11 | Extracellular Space | cytokine | 9MW3811 |
| -1.202 | 0.0314 | IL36G* | interleukin 36 gamma | Extracellular Space | cytokine | |
| -1.036 | 0.02 | PF4 | platelet factor 4 | Extracellular Space | cytokine | |
| -1.088 | 0.00583 | TNFSF13B* | TNF superfamily member 13b | Extracellular Space | cytokine | belimumab, tabalumab, telitacicept |
| 1.155 | 0.00366 | TNFSF15 | TNF superfamily member 15 | Extracellular Space | cytokine | |
| -1.81 | 0.0367 | WNT3A | Wnt family member 3A | Extracellular Space | cytokine | |

**Table 2. Ivacaftor altered cytokines in PANx cells (PANx_VX-770/ PANx_DMSO)**

**Asterick* represents downregulated genes in VX-770 treated PANx cell that were found to be elevated in PANx relative to the control ductal cells**
