## Supplementary Table for "Differential Expression and Microsystem Physiology Reveal Predominant and Drug Reversible CFTR-Related Defects in Idiopathic Pancreatitis"

### SUPPLEMENTARY MATERIAL

Includes supplementary Figure Legends, Material and Methods and supplementary references

### SUPPLEMENATRY FIGURE LEGENDS

#### Supplementary figure 1

**A.** Dot-plot showing top 10 genes corresponding to seven clusters with three major clusters identified as ductal mix (cluster 0,1,3,4), proliferating (cluster 2) and secretory/mucinous (cluster 5/6) based on gene ontology.

**B.** UMAP shows seven major ductal cell clusters in control and PANx cells which we broadly categorized into ductal mix, proliferating and secretory/mucninous populations.

#### Supplementary figure 2

**C.** Confocal image showing enlarged nuclie marked by DAPI signifying the presence of distinct cell population that we earlier identified to be mucinous.

**D.** Pancreatic lesion disease signature was obtained from ingenuity pathway analysis software (IPA, Qiagen©) and overlapped onto control, PANx-DMSO, PANx-VX-770 gene expression samples that showed enrichment for this signature in PANx and its reversibility using VX-770 based on differential expression gene (DEG) heat map.

**E.** Global transcriptomic data were mined to investigate the expression level of various genes that are known to be associated with cystic Intraductal papillary mucinous neoplasms (IPMN) and pancreatic ductal cell adenocarcinoma (PDAC). Each dot represents a single well containing the ductal organoids in the specified samples. The error bars represent S.E.M.

#### Supplementary figure 3

DEG map corresponding to the upregulated genes associated with **A.** Inflammation and specifically NF pathway and **B.** Lipid homeostasis and metabolism.

### MATERIALS AND METHODS

#### Generation and procurement of induced pluripotent stem cells (iPSCs)

The human iPSC lines were generated at Induced Pluripotent Stem Cell (iPSC) Core Facility, Cedars Sinai Biomanufacturing Center.The patient's IPSC line CS009iCP (PANx) was selected from the donor with idiopathic recurrent acute pancreatitis who was heterozygous for common pancreatitis CFTR variants c.1408 G>A and c.2562T>G with other two pancreatitis risk variants (heterozygous for variant in *SLC26A9* and homozygous for variant in *APOB*). The control line was selected from a healthy donor without any pancreatitis-related genetic risk variants and was age- and gender-matched.

#### iPSC maintenance and differentiation

Human induced pluripotent stem cells (iPSC) were cultured in feeder-free conditions using mTeSR™1medium (STEMCELL Technologies, #85850) supplemented with 10 µM Y-27632 on Matrigel-coated (Corning, #354277) plates. iPSCs were maintained at 37°C in 5% $CO_2$ until they reached 75%–95% confluency and were passaged every 4–6 days using Versene (Gibco, #15040066). For the differentiation process, iPSCs were dissociated using TrypLE select (Gibco #12563011) centrifuged (300 × g, 5 min), resuspended in mTeSR1 + Y-27632, and seeded at 20,000–30,000 cells per well (24-well plate). Cells were cultured for 24–48 hours to obtain 75%–95% confluency before initiating differentiation.

Directed differentiation of iPSC was performed using the previously published protocol with few modifications to generate pancreatic ductal-like organoids (PDLO) providing a population of pancreatic ductal cells(Breunig et al., 2021).

To induce mesendoderm differentiation, the medium was replaced with BE1a medium (MCDB131, 0.1% Fatty-acid-free BSA, 1.174 g/L Sodium bicarbonate, 0.8 g/L Glucose, 2 mM L-Glutamine) supplemented with 100 ng/mL Activin A and 2 µM CHIR99021 (Day 0). From Day 1 to Day 3, cells were cultured in BE1a containing 100 ng/mL Activin A and 5 ng/mL FGF2, with daily medium changes to induce definitive endoderm (DE). From Day 3 to Day 6, cells were cultured in BE1b medium (MCDB131, 0.5% Fatty-acid-free BSA, 1.174 g/L Sodium bicarbonate, 0.8 g/L Glucose, 2 mM L-Glutamine) supplemented with 50 ng/mL FGF10, 0.75 µM Dorsomorphin, and 3 ng/mL Wnt3a to promote gut tube endoderm (GTE) differentiation, with daily medium changes. Cells were cultured in BE3 medium (MCDB131, 2% Fatty-acid-free BSA, 1.754 g/L Sodium bicarbonate, 0.44 g/L Glucose, 2 mM L-Glutamine, 44 mg/L L-Ascorbic acid, 0.5X

ITS-X) containing 50 ng/mL FGF10, 200 nM LDN-193189, 0.25 µM SANT-1, 2 µM retinoic acid, and 16 mM glucose From Day 6 to Day 9, to induce pancreatic endoderm (PE). Between Day 9 and Day 13, differentiation into pancreatic progenitors (PPs) was achieved by culturing in BE3 medium supplemented with 100 ng/mL EGF, 200 nM LDN-193189, 10 mM Nicotinamide, 330 nM Indolactam V, and 16-mM glucose, with daily medium changes. At Day 13, PPs were harvested and resuspended in GFR-Matrigel (50 µL per 30,000 cells) before plating as 3D domes (50 µL per well, 24-well plate). After 10 minutes of incubation at 37°C, domes were overlaid with medium consisting of BE3 + 50 ng/mL EGF, 50 ng/mL FGF10, 50 ng/mL KGF, 50 nM MSC2530818, 10 mM Nicotinamide, 10 mM Y-27632, and 10 mM $ZnSO_4$(D13–D20). From Day 20 to Day 30, pancreatic trunk-like organoids (PTLOs) were further differentiated into pancreatic duct like organoids (PDLOs) using pancreatic ductal organoid (PDO) medium, was composed of Advanced DMEM/F12 with 1% penicillin/streptomycin, 1%FBS, 1%Glutamax and 10 mM HEPES, supplemented with 2% B27 supplement, 1% N2 supplement, 1 mM N-acetylcysteine, 0.5 nM Wnt3a, 100 ng/ml human R-spondin, 100 ng/ml human Noggin, 50 ng/ml recombinant human EGF, 100 ng/ml human FGF-10, 10 nM human gastrin and 1µM A83-0. Throughout differentiation, care was taken to avoid mechanical disruption of the thick matrigel layer during medium changes. Depending on the density, PDLOs were harvested between Day 27 and Day 30 and further expanded. Initial molecular characterization of PDLOs was performed at the initial passages: Passage 1 and Passage 2.

**Human pancreatic ductal organoid culture**

Human pancreatic ductal organoids were cultured as described previously(Breunig *et al.*, 2021) with some modifications. Pancreatic ductal cells were suspended in GFR-Matrigel (50,000 cells per 50 µL) plated as 3D domes (50 µL dome per well (24-well Thermofisher# 142475)) and overlaid with 500 µL PDO medium. Medium change was done every 2-3 days and organoids were passaged every 5-7 days using TrypLE Select.

**Human pancreatic ductal monolayer culture**

Pancreatic ductal organoids were dissociated into single cells using TrypLE Select, and cultured ($1X10^5$ cells) on Matrigel (Corning, #354230) (1mg/6ml) precoated transwell cell culture inserts (24-well, Transwell, Costar #3470-Clear, 0.4 mM pore size) until they formed a polarized monolayer. Media change was performed every 2-3 days (Apical 400 µL, basolateral 600 µL).

**Short Circuit Current measurements**

Pancreatic ductal monolayers were obtained on transwell cell culture inserts. Once the transepithelial electrical resistance (TEER) exceeded 1000Ω·cm² (measured using an EVOM system, WPI), the inserts were transferred to an Ussing chamber inserts, as previously described (Clarke, 2009). Both the apical and basolateral chambers were filled with Krebs-Ringer's buffer solution, and the temperature was maintained at 37°C using a circulating water bath. CFTR function was assessed by measuring changes in short-circuit current ($I_{sc}$) in response to CFTR agonist- Forskolin (FSK, 4 µM), CFTR inhibitor- CFTRinh-172 (10 µM), and CFTR potentiator- Ivacaftor (VX-770, 4 µM).

**MUC5AC ELISA**

Pancreatic ductal monolayers cultured on Transwell inserts were treated with DMSO or the CFTR potentiator Ivacaftor (VX-770, 4 µM) for 48 hours**.** Following treatment, the inserts were treated with 1 mM dithiothreitol (DTT) to break disulfide bonds within the mucins and enable mucin depolymerization and detachment from epithelial surface. After 1 h of DTT treatment, 100 µL of apical media was collected for measuring secreted MUC5AC using the Invitrogen ELISA Kit (#EEL062) following the manufacturer's instructions.

**FSK induced fluid secretion measurements**

Pancreatic ductal organoids were split into single cells using TrypLE and about 10,000 cells were plated as 50 µl of 50% Matrigel-media suspension per well in a 96-well clear bottom plate. The cells were allowed to form organoids for 3-4 days and were subjected to FSK-induced fluid secretion assay under various treatment combination with FSK alone or FSK plus various phosphodiesterase inhibitors. FSK-induced swelling (FIS) was monitored over 72 h using time-lapse microscopy (Lionheart FX, Agilent BioTek). FIS was quantitated as total organoid area using the FIJI, ImageJ software.

**Whole cell patch clamp recording**

For whole-cell patch-clamp recordings, composition of pipette and bath solution performed from the experiments were based on our previously published experiments with pipettes resistances ranging between 3 - 6 MΩ (Lee et al., 2024) . The current traces and I/V curves of CFTR responses were consecutively recorded with a 1 s voltage ramp of ± 100 mV applied every 10 s: hold at Vm = - 60 mV and filtered at 1 kHz and sampled at 50 Hz. To enable robust CFTR, we used intracellular cAMP (100 μM) in addition to FSK and applied through the pipette and the CFTR specific currents were confirmed using CFTRinh-172 (20 μM).

**Pancreatic Tissue Dissociation and Acinar Cell Isolation**

Human pancreatic tissue was obtained from Pancreas Care Center, Nationwide Children's Hospital, Columbus, OH. The pancreatic tissue was manually dissected to remove fat, connective tissue, and visible blood vessels. The tissue was then minced into small fragments (~0.5–1 $mm^3$) using sterile surgical scissors. The minced tissue was enzymatically digested in HBSS containing $Ca^{2+}$ and $Mg^{2+}$, supplemented with 0.5 mg/mL Collagenase P (Roche), 0.5 mg/mL Dispase II (Sigma-Aldrich), 0.1 mg/mL Trypsin Inhibitor, and 0.1 mg/mL DNase I (Sigma-Aldrich). Digestion was carried out at 37°C for 1–2 hours with continuous agitation in an orbital shaker. Following enzymatic digestion, the tissue was mechanically dissociated by gentle trituration using a 10 ml serological pipette and cell suspension was passed through a 70 μm cell strainer to remove undigested tissue and the filtrate was centrifuged at 300 × g for 5 minutes at 4°C to pellet the acinar-rich fraction. The cell suspension was washed twice with PA medium consisting of Advanced DMEM/F12 supplemented with 1% penicillin/ streptomycin, 1% FBS, 1% GlutaMAX, and 10 mM HEPES, then resuspended in the same medium for culture.

**Modeling the communication between hyper viscous ductal fluid and acinar cells using 3D μ-Slide chip**

For acinar cell culture**,** pancreatic acinar (PA) medium was used, consisting of Advanced DMEM/F12 supplemented with 1% penicillin/streptomycin, 1% FBS, 1% GlutaMAX, and 10 mM HEPES**.** Isolated acinar clusters were suspended in a Matrigel and PA medium mixture (3:2 ratio)**.** A 30 μL cell suspension was seeded into wells on one side of a three-channel μ-Slide (ibidi #80376), while the opposite wells connected by a narrow channel were filled with PA medium. The μ-Slide chip was then sealed**,** and the channels and reservoirs were filled with PA medium to prevent the formation of air bubbles**.** Media was refreshed every 2–3 days**.** Meanwhile, pancreatic ductal cells were cultured on Transwell inserts to form a ductal monolayer. Once the tight monolayer was formed at 1000Ω·$cm^2$ in < 7 days, the cells were maintained on the inserts for additional 48 h without change of media allowing for the collection of pancreatic ductal fluid within the apical media. After 5–7 days of acinar culture in the three-channel μ-Slide, the channel without cells and only PA medium was replaced with 30 μL of pancreatic ductal media collected from apical side of the Transwell-cultured ductal cells. To modify the viscosity of the pancreatic ductal media, it was mixed with 10% glycerol while keep the effective ductal fluid composition same in two treatment conditions i.e., with and without glycerol. Acinar cells were treated with the ductal fluid for 24 hours, after which the entire medium was collected for testing amylase secretion and lactate dehydrogenase activity.

**Amylase activity to measure acinar cell function on the microchamber device**

After 24 hours of treatment with the ductal medium with and without glycerol**,** 100 µL of the medium was collected from each channel from the 3D µ-Slide chip**.** The amylase activity of each sample was then assessed using the Abcam Amylase Activity Colorimetric Assay Kit (#ab102523), following the manufacturer's instructions. Amylase activity was quantitated as nmoles/min.

**Lactate dehydrogenase activity to measure acinar cell viability**

After 24 hours of treatment with glycerol-supplemented ductal medium, 1µL of pancreatic acinar supernatant was collected from each channel of the 3D µ-Slide chip and mixed with 49 µL of LDH Storage Buffer (200mM Tris-HCl (pH 7.3), 10% Glycerol, 1% BSA) to prepare samples. The LDH activity of each sample was then assessed calorimetrically using LDH-Glo™ Cytotoxicity Assay (Promega Cat#J2380), following the manufacturer's instructions.

**Immunofluorescence Staining of Pancreatic Ductal Cell Monolayers**

Pancreatic ductal cell monolayers were fixed using 3.7% paraformaldehyde (Sigma, Cat#252549) for 15 minutes at room temperature (RT), then permeabilized for 30 minutes at RT using 10X Permeabilization Buffer (Invitrogen, Cat#00-8333-56) that was diluted 1:10 in double-distilled water. Following permeabilization, samples were blocked with 2.5% normal goat serum (Vector Laboratories, Cat#S-1012-50) for 2 hours at RT. The cells were then incubated overnight at 4°C with primary antibodies, anti-CFTR R1104 (Eric Sorscher lab, CF Center, University of Alabama, Birmingham, AL, USA [presently, Emory University, Atlanta, GA,USA]), anti-ZO-1 (BD Biosciences; #610967), anti-KRT 19 (Invitrogen; #MA5-12663), anti-E cadherin (Cell Signaling Technology; #3195), diluted 1:100 in antibody diluent buffer (Epredia, Cat#TA-125-ADQ).The following day, samples were washed for 5 minutes three times with PBS + 0.05% Tween-20 (Sigma, Cat#P9416-100ML) and incubated with the secondary antibodies, (Invitrogen; Alexa Fluor 488 or 568 diluted 1:500 in in antibody diluent buffer**,** for 2 hours at RT. After secondary antibody incubation, samples were washed twice for 5 minutes each with PBS + 0.05% Tween-20, then incubated with DAPI solution (ThermoFisher, Cat#62248), diluted 1:1000 in PBS, for 5 minutes at RT, followed by a final third wash. Finally, samples were mounted using Vectashield antifade mounting medium (Vector Laboratories, Cat#H-1000), and imaging was performed using Nikon AX R NSPARC Confocal Microscope.

**Immunofluorescence Staining of Pancreatic Ductal Cell Monolayers for Mucin**

Pancreatic ductal cell monolayers cultured on Transwell inserts were treated with either DMSO (vehicle control) or the CFTR potentiator Ivacaftor (VX-770, 4 µM; Selleckchem) for 48 hours. Following treatment, cells were fixed with 3.7% paraformaldehyde (Sigma, Cat# 252549) for 15 minutes at room temperature (RT), then permeabilized for 30 minutes at RT using 1X Permeabilization Buffer (Invitrogen, Cat# 00-8333-56), prepared by diluting the 10X stock 1:10 in double-distilled water. After permeabilization, the Transwell membranes were carefully excised from the inserts and placed onto glass microscope slides. Samples were blocked in 5% bovine serum albumin (BSA) in PBS containing 0.05% Tween-20 (PBST) for 2 hours at RT. This was followed by overnight incubation at 4°C with 10 µg/mL of the biotinylated Mucin-specific probe, StcE (Sigma-Aldrich, Cat# SAE0212), diluted in antibody diluent buffer (Epredia, Cat#TA-125-ADQ).The next day, samples were washed three times for 10 minutes each in PBST, then incubated with NeutrAvidin (5 µg/mL; ThermoFisher, Cat# 84606) for 60 minutes at RT. After two additional 10-minute washes in PBST, nuclei were counterstained with DAPI (1 µg/mL; ThermoFisher, Cat# 62248) in PBST for 5 minutes at RT, followed by a final wash step. Finally, samples were mounted using VECTASHIELD Antifade Mounting Medium (Vector Laboratories, Cat# H-1000) and imaged using Nikon AX R NSPARC Confocal Microscope.

**Immunohistochemistry**

Pancreatic ductal cell monolayers grown on Transwell inserts were fixed with 3.7% paraformaldehyde for 15 minutes at room temperature (RT), then washed three times with PBS. After fixation, the Transwell membrane was carefully separated from the insert by cutting around the edge, ensuring minimal disruption to the monolayer. The excised membrane was then embedded (cell side up) in a 3% agar drop, maintained at approximately 50–55°C to prevent thermal damage. The prepared samples were submerged in PBS and sent to the Cedars-Sinai Biobank for paraffin embedding, sectioning, and histological staining for Hematoxylin and eosin (H&E) for general tissue morphology and Alcian Blue-Periodic Acid-Schiff (AB-PAS) staining for detecting acidic and neutral mucins.

**RNA Extraction for Bulk-RNA sequencing**

Total RNA extraction and poly(A)-enriched directional RNA sequencing were carried out by the Genomics, Epigenomics, and Sequencing Core Facility at the University of Cincinnati(Qiu et al., 2023; Reigle et al., 2021).

In brief, RNA integrity was assessed using an Agilent Bioanalyzer (Santa Clara, CA), confirming high-quality samples with RIN scores above 9. Library preparation was performed using the

NEBNext Ultra II Directional RNA Library Prep Kit for poly(A) RNA. Following library quality assessment via Bioanalyzer and quantification using the NEBNext Library Quant Kit (New England BioLabs), libraries with unique indices were pooled in appropriate ratios and sequenced on the Illumina NextSeq 550 platform (San Diego, CA). Subsequent bioinformatic analysis was conducted through the BaseSpace SEQUENCE HUB using the RNA-Seq Alignment app v2.0.2 and the RNA-Seq Differential Expression app v1.0.1. STAR was utilized for read alignment, Salmon for transcript quantification (reported in TPM), and DESeq2 for identifying differentially expressed genes. Bulk RNA-seq data visualization was carried out using custom R scripts. Heatmaps was generated to explore sample clustering, expression patterns, and differential gene expression using Morpheus.

**Single-Cell RNA Sequencing and Data Processing**

Single-cell suspensions were prepared from both PANx and control samples and single-cell libraries were generated according to the manufacturer's protocol (10x Genomics, Single Cell 3′ Reagent Kits v3; https://www.10xgenomics.com/products/single-cell). Cellular suspensions were loaded onto a Chromium Controller (10x Genomics) to generate single-cell Gel Bead-in-Emulsions (GEMs). Reverse transcription (RT) was performed within GEMs using a Veriti 96-well thermal cycler (Thermo Fisher Scientific). Following RT, GEMs were broken, and the barcoded complementary DNA (cDNA) was recovered and purified. Size selection was carried out using the SPRIselect Reagent Kit (Beckman Coulter). Indexed sequencing libraries were prepared using the Chromium Single Cell 3′ Library Kit (10x Genomics), which includes enzymatic fragmentation, end-repair, A-tailing, adapter ligation, ligation cleanup, sample indexing PCR, and final library cleanup. Prepared libraries were sequenced on an Illumina NextSeq X Plus platform at Novogene (https://www.novogene.com/).Single-cell RNA sequencing was performed to characterize transcriptomic profiles in ductal epithelial cells from PANx and control conditions. Approximately 11,000 cells were captured from the PANx group and 8,000 cells from the control group. Sequencing reads were aligned to the reference genome using Cell Ranger (v6.1.2) to generate gene-barcode matrices for each sample. All downstream analyses were conducted using R version 4.4.1 and the Seurat package (https://satijalab.org/seurat/) (v5.0.1). Initial quality control (QC) steps included filtering out cells with fewer than 200 detected genes or more than 10% mitochondrial gene content. After QC filtering, 6,475 cells from the PANX group and 5,196 cells from the control group were retained for downstream analysis. Doublet detection and removal were performed using the scDblFinder package (v1.16.0) integrated with Bioconductor workflows. Doublets were identified and excluded prior to normalization and clustering. Filtered datasets were processed using Seurat's standard workflow. Each dataset was log-normalized and the top 2,000 variable features were identified. The datasets were then integrated using Seurat's reciprocal PCA-based integration pipeline. Following integration, dimensionality reduction was performed using principal component analysis (PCA), and the top 15 PCs were used for UMAP embedding and graph-based clustering. Clusters were identified and with a resolution of 0.5. Expression patterns of key genes, including mucin family genes, were visualized on the UMAP, revealing their enrichment in a distinct mucinous ductal cell population.

**Statistical Analysis**

All data are presented as mean ± standard deviation (SD) and were obtained from a minimum of two independent experimental replicates. Statistical analyses were conducted using GraphPad Prism software (version 10). For pairwise comparisons, an unpaired Student's *t*-test was used. When comparing three or more groups, either one-way or two-way ANOVA was performed, depending on the number of independent variables, followed by Tukey's multiple comparison test. A *P*-value of less than 0.05 was considered statistically significant.

**REAGENT AND RESOURCE TABLE**

| REAGENT or RESOURCE | SOURCE | IDENTIFIER |
|---|---|---|
| Antibodies | | |
| Alexa Fluor 488 Goat anti-Rabbit IgG antibody | Invitrogen | A-11008 |
| Alexa Fluor 568 Goat anti-mouse IgG antibody | Invitrogen | A-11004 |
| Alexa Fluor 568 Donkey anti-Sheep IgG antibody | Invitrogen | A-21099 |
| Alexa Fluor 568 Goat anti-Rabbit IgG antibody | Abcam | ab175471 |
| E-Cadherin Rabbit | Cell Signaling | 3195 |
| Mouse anti-Human KRT 19 | Invitrogen | MA5-12663 |
| Mouse anti-Human ZO-1 | BD Biosciences | 610967 |
| Mucin Probe, StcE | Sigma | Mucin Probe, StcE |
| Pancreatic amylase alpha antibody | Novus Biologicals | NB10066388 |
| Chemicals, peptides, and recombinant proteins | | |
| (-)-Indolactam V | STEMCELL | 72314 |
| 2.5% normal goat serum | Vector Laboratories | S-1012-50 |
| 8-Br-cAMP | Sigma | B7880-25MG |
| Activin A | R&D systems | 338-AC-010 |
| Advanced DMEM/F12 | Gibco | 12634028 |
| Antibody diluent buffer | Epredia | TA125ADQ |
| B-27 suppliments 50X | Thermo Fisher Scientific | 17504044 |
| CFTRinh-172 | Sigma | C2992 |
| Cilostazol | Tocris | 1692 |
| Collagenase/Dispase® Roche | Sigma | 11097113001 |
| Corning® Matrigel® Basement Membrane Matrix, LDEV-free | Corning | 354234 |
| Corning® Matrigel® Growth Factor Reduced (GFR) | Corning | 354230 |
| Corning® Matrigel® hESC-Qualified Matrix | Sigma | CLS354277 |
| Corning® Matrigel® hESC-Qualified Matrix, LDEV-free | Corning | 354277 |

| | | |
|---|---|---|
| DAPI solution | Thermo Fisher Scientific | 62248 |
| DMSO | Merck | D2650 |
| DNase 1 solution | stemcell | 7900 |
| Dorsomorphin | stemgent | 04-0024 |
| DTT (dithiothreitol) | Thermo Fisher Scientific | R0861 |
| FA free BSA (catalog item) | Sigma | A7030-10G |
| Fatty acid free BSA | Sigma | A7030-10G |
| FBS | Sigma | 12103C |
| Glucose solution (sterile) | Thermo Fisher Scientific | A2494001 |
| Forskolin | Sigma Aldrich | F3917-10MG |
| Glucose solution (sterile) | Thermo Fisher Scientific | A2494001 |
| Glycerol, Molecular Biology Grade | Thermo Fisher Scientific | J61059.K2 |
| HEPES(1M) | Thermo Fisher Scientific | 15630106 |
| Human FGF 10 | R&D systems | 345-FG-025 |
| Human Gastrin I | R&D systems | SKUG9145-.1MG |
| Insulin-Transferrin-Selenium-Ethanolamine | Thermo Fisher Scientific | 51500056 |
| Ivacaftor (VX770) | Selleckchem | S1144 |
| L-Glutamine | Sigma | 25030081 |
| LDN193189 hydrochloride | Sigma | SML0559-5MG |
| LDN193189 hydrochloride | Sigma | SML0559-5MG |
| MCDB 131 Medium | Thermo Fisher Scientific | 10372019 |
| MSC2530818 | Selleckchem.com | S8387 |
| mTeSR™1medium | STEMCELL | 85850 |
| N-2 Supplement | Fisher Scientific | 17502048 |
| N-Acetylcysteine | R&D systems | 7874/100 |
| Neutravidin | Thermo Fisher | 84606 |
| Nicotinamide | Sigma | N0636 |
| Paraformaldehyde 37% | Sigma | 252549 |
| Permeabilization Buffer | Invitrogen | 00-8333-56 |

| | | |
|---|---|---|
| Penicillin-Streptomycin | Gibco | 15-140-163 |
| Recombinant Human EGF Protein, CF | R&D systems | 236-EG-200 |
| Recombinant Human FGF-4 Protein | R&D systems | 235-F4-01M |
| Recombinant human FGF2 | R&D systems | 233-FB-025 |
| Recombinant Human KGF (FGF-7) | Sigma | SRP3100 |
| Recombinant Human Noggin Protein with carrier | R&D systems | 6057-NG-100 |
| Recombinant Human R-Spondin 1 Protein | R&D systems | 4645-RS-025 |
| Recombinant Human Wnt-3a Protein | R&D systems | 5036-WN-010/CF |
| Retinoic Acid | Sigma | R2625 |
| Rolipram | Tocris (Bio-Techne) | 0905 |
| RPMI 1640 Medium, GlutaMAX™ Supplement, 500 mL | Thermo Fisher Scientific | |
| SANT-1 | Tocris (Bio-Techne) | 1974 |
| Sodium bicarbonate | Sigma | S8875 |
| soybean trypsin inhibitor | Thermo Fisher Scientific | 17075029 |
| Tris-Acetate-EDTA(TAE), 50X Solution | Fisher Scientific | BP1332500 |
| TrypLE™ Select Enzyme (1X), no phenol red | Thermo Fisher Scientific | 12563011 |
| Tween-20 | Sigma | P9416 |
| Vectashield antifade mounting medium | Vector Laboratories | H-1000 |
| Versene solution | Gibco | 15040066 |
| Y-27632 dihydrochloride | Tocris (Bio-Techne) | 1254 |
| Zaprinast | Tocris (Bio-Techne) | 0947 |
| ZnSO4 | Sigma | Z1001 |
| Critical commercial assays | | |
| LDH-Glo™ Cytotoxicity Assay | Promega | J2380 |
| MUC5AC ELISA Kit | Invitrogen | EEL062 |
| Experimental models: Cell lines | | |
| CS009iCP | Cedars-Sinai Biomanufacturing Center | n/a |

| | | |
|---|---|---|
| CS2GW3iCTR | Cedars-Sinai Biomanufacturing Cnter | n/a |
| Other | | |
| Costar® 6.5 mm Transwell®, 0.4 µm Pore Polyester Membrane Inserts | Stem cell technologies | 38024 |
| Nunclon treated 24 well plates | Thermo Fisher Scientific | 142475 |
| µ-Slide III 3D Perfusion | ibidi | 80376 |
