## Supplementary figures and images for "Differential Expression and Microsystem Physiology Reveal Predominant and Drug Reversible CFTR-Related Defects in Idiopathic Pancreatitis"

### Supplementary figures

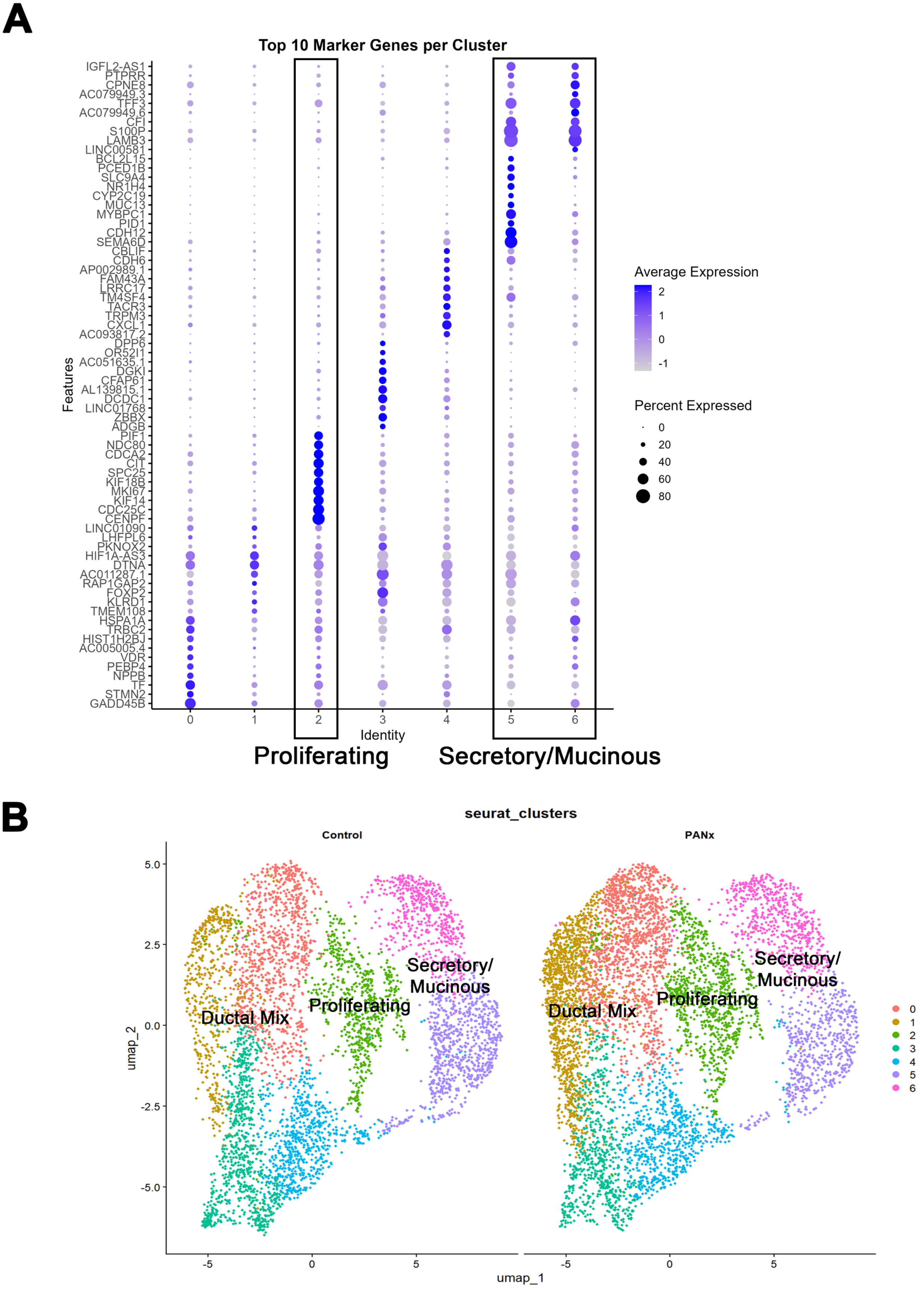


**Supplementary Figure 1**

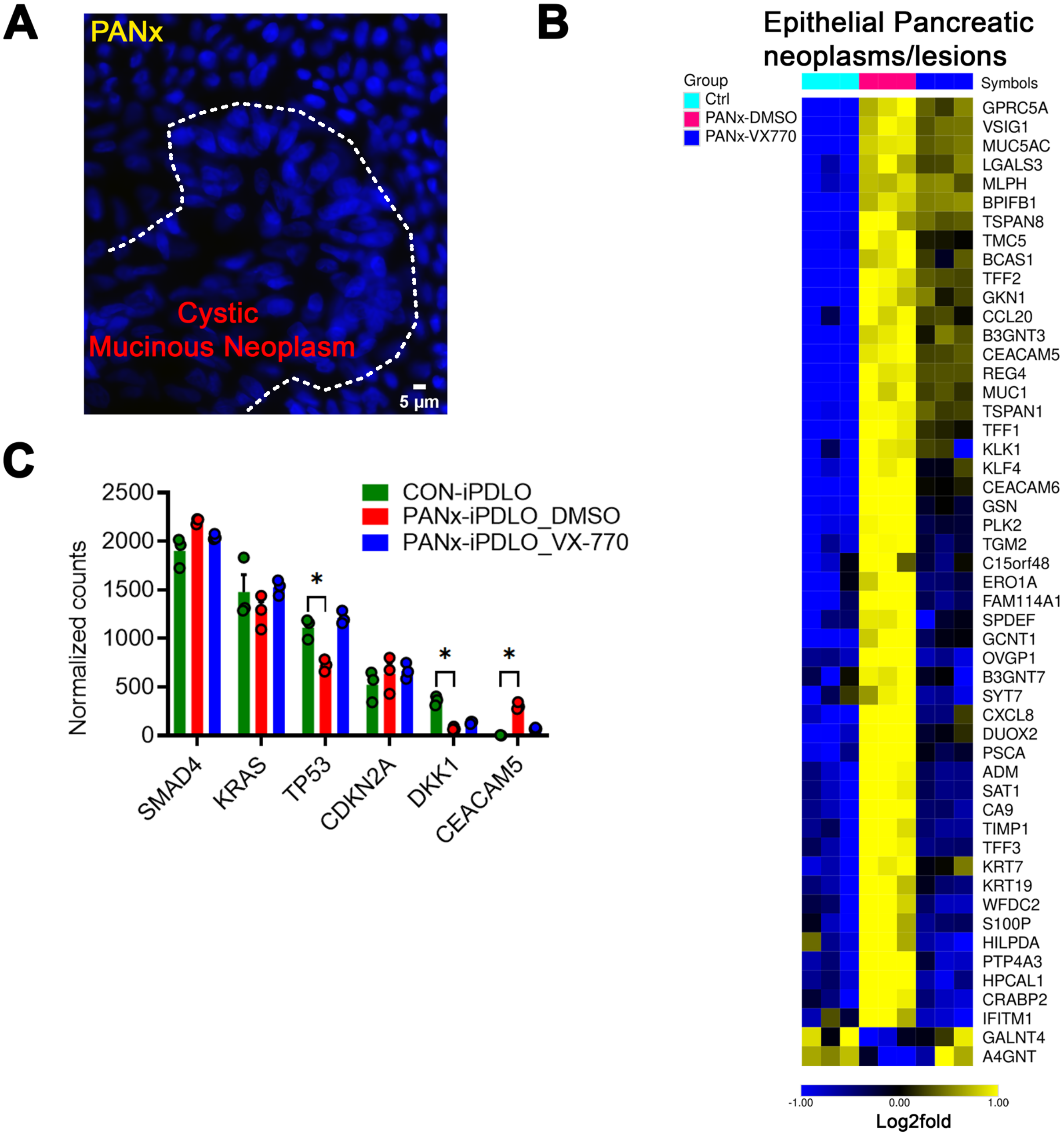


**Supplementary Figure 2**

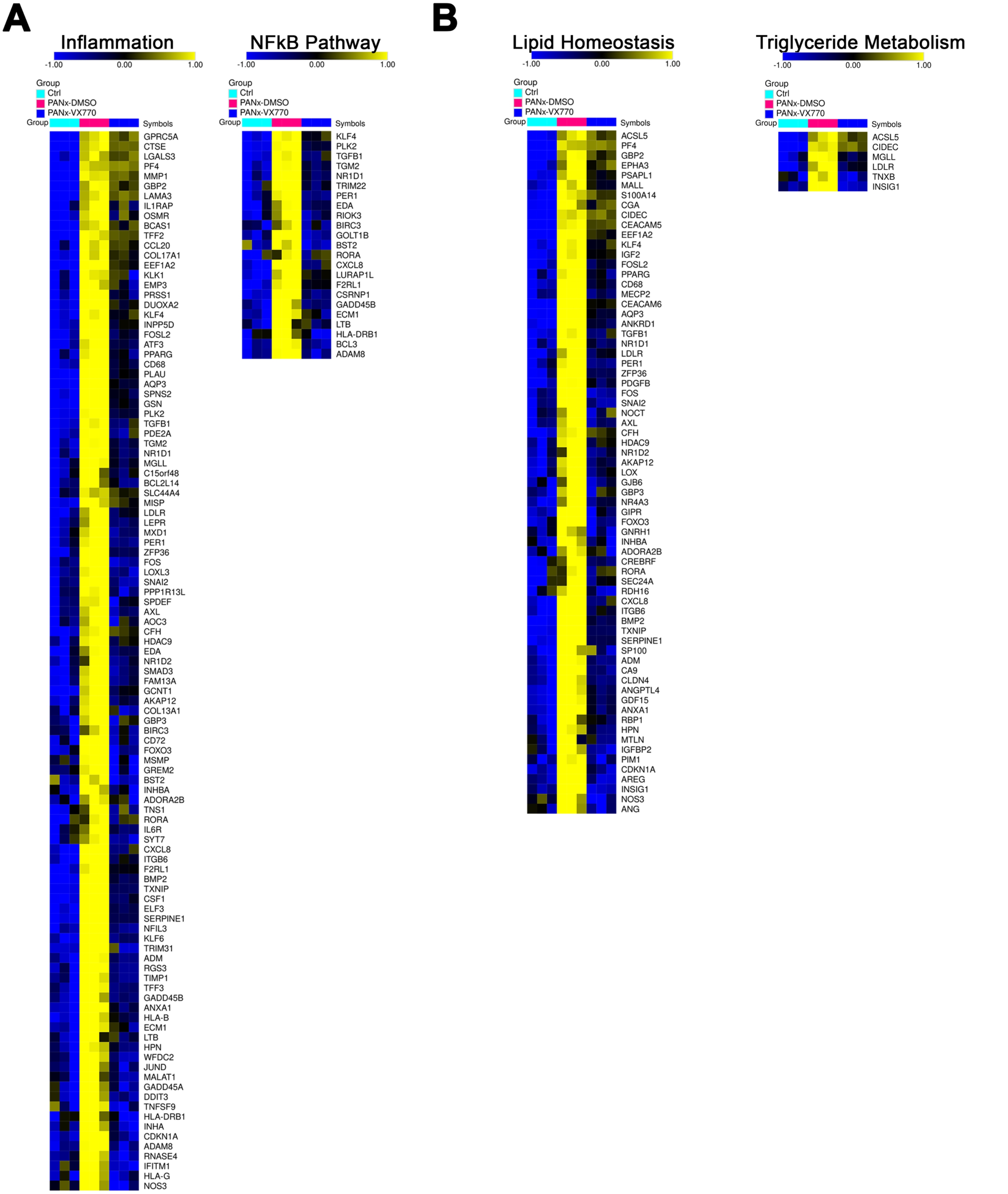


**Supplementary Figure 3**
